# Out of Place: Three Eurasian Taxa Shed Light on Origin of Thermopsideae in North America

**DOI:** 10.1101/2023.10.21.563365

**Authors:** Todd A. Farmer, Robert K. Jansen

## Abstract

The North American Thermopsideae, a monophyletic group comprising the North American endemic *Baptisia*, and the paraphyletic Eurasian–North American disjunct *Thermopsis*, is nested within the Sophoreae (Fabaceae: Papilionoideae). Previous phylogenetic studies have identified two East Asian taxa within the North American Thermopsideae, suggesting two independent dispersal events between North America–East Asia. More recent studies have also placed a third taxon, *Vuralia turcica*, an endemic species from Turkey, among the North American Thermopsideae. The presence of three geographically distant Eurasian taxa within a relatively young clade of North American origin is perplexing, and the biogeographic implications of this observation are not clear. To investigate this matter, 1540 low-copy nuclear genes and complete plastomes were obtained from 36 taxa across the core genistoids, including 26 newly sequenced taxa. Nuclear and plastome based maximum likelihood (ML) and ASTRAL analyses were conducted based on varying degrees of taxon coverage and read mapping consensus threshold values. Additional analyses were performed to estimate divergence times and to reconstruct biogeographic history. The results strongly support a relictual Old World clade, presently composed of *V. turcica* and *T. chinensis*, which diverged from the ancestor of the North American lineage during the mid to late Miocene. A single and recent North America–East Asia dispersal involving *T. lupinoides* is reported. Furthermore, the traditional inclusion of the genus *Ammopiptanthus* among Thermopsideae, and the recent recircumscription of *T. turcica* to the monotypic *Vuralia*, are not supported. A relatively high degree of cytonuclear discordance is reported within each sub-clade of the North American Thermopsideae. This finding is likely attributable to the high degree of interspecific hybridization reported within these groups and raises the need for more rigorous genome-scale testing to disentangle their complex phylogenies.

**Subjects:** Biodiversity, Biogeography, Evolutionary Studies, Genetics, Plant Science

## Introduction

Thermopsideae (Yakovlev 1972) are a relatively small tribe in Fabaceae (Papilionoideae) that traditionally includes six genera, *Ammopiptanthus* S.H. Cheng, *Anagyrus* L., *Baptisia* Vent, *Pickeringia* Nutt., *Piptanthus* Sweet, and *Thermopsis* R.Br. Though the genera comprising Thermopsideae were widely accepted by early works, more recent phylogenetic analyses have resulted in major revisions to the group, including the transfer of *Pickeringia* into the *Cladrastis*-*Styphnolobium* clade, and the addition of a new monotypic genus, *Vuralia* Uysal & Ertuğrul (Turner 1981; Kit et al., 1982; Wojciechowski et al., 2004; Uysal et al., 2014). Additionally, the placement of *Ammopiptanthus* within Thermopsideae has been problematic. In Wang et al. (2006) *Sophora* formed a clade with the core Thermopsideae, sister to *Ammopiptanthus*, and in Cardoso et al. (2013) *Sophora* formed a clade with *Ammopiptanthus*, sister to the core Thermopsideae. These findings resulted in the merging of Thermopsideae into Sophoreae *sensu* Cardoso, making the group a monophyletic member of the core genistoids. Although this interpretation was not supported by Zhang et al. (2015), who placed *Ammopiptanthus* in Thermopsideae, distant from *Sophora*, a number of subsequent phylogenetic studies have since supported Sophoreae *sensu* Cardoso, which includes a strongly supported thermopsoid clade comprising ca. 45 species and five genera – the Sino-Himalayan *Piptanthus*, the Micronesian-Mediterranean *Anagyrus*, the Turkish endemic *Vuralia*, the Eurasian-North American disjunct *Thermopsis*, and the North American endemic *Baptisia* (Zhang et al., 2015; Shi et al., 2017; Choi et al., 2022)

Previous studies indicate a monophyletic North American Thermopsideae, composed of two sister clades, *Baptisia,* with 17 species occurring in the grasslands and coastal plains of the Eastern and Southeastern regions of the United States, and the paraphyletic *Thermopsis*, with seven species occurring in mountainous regions of Western United States, and three species occurring in the southeastern Appalachian foothills (Mendenhall 1994; Chen et al., 1994; Sa et al., 2000; Turner 2006; Wang et al., 2006; Cardoso et al., 2012; Cardoso et al., 2013; Uysal et al., 2014; Zhang et al., 2015; Shi et al., 2017; Choi et al., 2022). Based on morphology, both Czefranova (1970) and Sa et al. (2000) grouped the only two northeastern Asian endemic species of *Thermopsis*, *T. lupinoides* Link (synonym: *T. fabacea* DC.) and *T. chinensis* Benth. ex S. Moore, with the North American element, to form sections *Thermia*, and *Archithermopsis*, respectively. Numerous subsequent phylogenetic studies have supported this interpretation, placing *T. lupinoides* within the North American *Thermopsis* clade and *T. chinensis* sister to either the North American *Thermopsis* or *Baptisia* (Wang et al., 2006; Zhang et al., 2015). More recently, the plastome markers *matK, rbcL, trnL-trnF* and *psbA-trnH*, as well as the nuclear ribosomal DNA internal transcribed spacers (ITS 1 and 2) analyses of Shi et al. (2017) placed a third Asian taxon, *Vuralia turcica* (Kit Tan, Vural & Küçük.) Uysal & Ertuğrul (which was excluded from the previous analyses) within the North American *Thermopsis* clade. The ITS marker placed this taxon sister to *T. chinensis* and distant from *T. lupinoides*. These findings have been interpreted as the result of multiple independent dispersals between East Asia and North America, the first involving the initial colonization of North America, driven by the Qinghai-Tibetan Plateau uplift (QTP), followed by at least two subsequent dispersals back into Asia by the ancestors of *T. lupinoides, T. chinensis*, and *V. turcica,* (Zhang et al., 2015; Shi et al., 2017).

Given the rarity of East Asia–North America disjunction among papilionoid legumes, multiple independent occurrences of this phenomenon within a relatively small and young clade is perplexing, and although many molecular phylogenetic studies concerning Thermopsideae have been conducted, they have traditionally relied on a limited number of markers, and many have excluded *Ammopiptanthus*, *Vuralia*, and most of the North American Thermopsideae. A single study has included both *V. turcica* and *T. chinensis* – two species that may be of great importance to understanding the biogeographic history of the North American Thermopsideae (Mendenhall, 1994; Käss & Wink, 1995, 1996, 1997a, b; Doyle et al., 1997, 2000; Crisp et al., 2000; Kajita et al., 2001; Pennington et al., 2001; Heenan et al., 2004; Wojciechowski et al., 2004; Wang et al., 2006; Uysal et al., 2014, Zhang et al., 2015, Shi et al., 2017).

Modern advances in molecular sequencing technology have greatly improved the feasibility of obtaining high-throughput molecular sequence data, which compared to traditional phylogenetic markers have been shown to enhance evolutionary signal. The goal of the present study was to generate high throughput nuclear and plastid sequence data to: (1) establish a core backbone phylogeny for Thermopsideae in a core genistoids framework; (2) test the monophyly of Thermopsideae, particularly with respect to the placement of *Ammopiptanthus*; (3) test the monophyly of the North American Thermopsideae; and (4) resolve the phylogenomic placement and biogeographic significance of *V. turcica*, *T. chinensis*, and *T. lupinoides*.

## Materials and Methods

### Taxon Sampling

The taxon sampling strategy was designed to include all major clades and subclades of Thermopsideae, with an emphasis on taxa in need of additional investigation (Table 1). Of the 36 sampled taxa, 26 were newly sequenced. Of these, 10 were sequenced from fresh leaf material collected within native ranges, six from leaf material grown from seeds obtained from the United States Department of Agriculture (USDA), three from silica dried leaf material obtained from the living collection at the Missouri Botanical Garden, and seven from dried herbarium specimens. All fresh leaf samples were flash-frozen in liquid nitrogen and stored in an ultra-low freezer (-80°C) prior to DNA isolation. Vouchers for the relevant taxa were deposited in the Billie L. Turner Plant Resources Center at the University of Texas at Austin (TEX/LL; Table 1). The sampled taxa included a single accession of *Vuralia*, all North American *Thermopsis* (10 species; Chen et al., 1994), two members of each of the five morphological clades of *Baptisia* (Turner 2006), six members of the Asian *Thermopsis*, one member of *Ammopiptanthus*, one member of *Piptanthus*, and outgroups from Sophoreae (*Euchresta tubulosa* Dunn) and the core genistoids group (*Crotalaria albida* B. Heyne ex Roth and *Lupinus albus* L.; Choi et al., 2022). Except for the three Eurasian species that are allied to the North American Thermopsideae, the Eurasian taxa were primarily selected based on GenBank availability, which excluded *Anagyrus* at the time of sampling.

**Table 1:** Terminal taxa used in the phylogenomic analyses, including identifiers of vouchers (herbarium acronym, collectors name, and collection number), collection location, GenBank accession number, and mean plastome depth of coverage, when applicable. N/A = not available.

| ID | Taxon Name | Voucher Location | Collection Location | Accession Number | Mean Plastome Depth |
| --- | --- | --- | --- | --- | --- |
| 1G | *Thermopsis lanceolata* | TEX - Farmer 318 | Seed - USDA PI 678914 | N/A | 4416.7 |
| 2G | *Thermopsis dahurica* | TEX - Farmer 319 | Seed - USDA W6 19718 | N/A | 6384 |
| 3G | *Thermopsis montana*<br>var. *ovata* | TEX - Farmer 320 | Seed - USDA W6 10173 | N/A | 1813.3 |
| 4G | *Thermopsis davaricarpa* | TEX - Farmer 321 | Seed - USDA 90-0309 | N/A | 2362.5 |
| 6G | *Thermopsis rhombifolia* | TEX - Farmer 323 | Seed - USDA 90-0005 | N/A | 1224.8 |
| 7G | *Thermopsis gracilis* | TEX - Farmer 324 | Seed - USDA W6 54284 | N/A | 2711.4 |
| 12X | *Baptisia simplicifolia* | TEX - Farmer 200 | Liberty Co., FL | N/A | 568.7 |
| 18X | *Baptisia calycosa* | TEX - Farmer 206 | St. Johns Co., FL | N/A | 806 |
| 22X | *Baptisia lanceolata*<br>var. *lanceolata* | TEX - Farmer 210 | Nassau Co., FL | N/A | 1081.2 |
| 24X | *Baptisia arachnifera* | TEX - Farmer 212 | Brantley Co., GA | N/A | 1022.3 |
| 31X | *Baptisia hirsuta* | TEX - Farmer 219 | Walton Co., FL | N/A | 469.8 |
| 25B | *Baptisia nuttalliana* | TEX - Farmer 25 | Angelina Co., TX | N/A | 1193.7 |
| 31R | *Thermopsis mollis* | TEX - Farmer 299 | Cherokee Co., SC | N/A | 1720.9 |
| 33R | *Thermopsis villosa* | TEX - Farmer 301 | Henderson Co., NC | N/A | 2127.5 |
| 35R | *Thermopsis fraxinifolia* | TEX - Farmer 303 | Monroe Co., TN | N/A | 1285.7 |
| 33T | *Thermopsis montana* | TEX - Wojciechowski<br>2300 | Coconino Co., AZ | N/A | 1722.7 |
| H12 | *Baptisia cinerea* | TEX - Mendenhall 372 | Chesterfield, SC | N/A | 316 |
| H13 | *Baptisia alba* | TEX - T. Wayne Barger | Jackson Co., SC | N/A | 253.3 |
| H19 | *Thermopsis turcica* | RSA-POM - Kit Tan and<br>Kucukoduk 19-A | Aksehir, Turkey | N/A | 982.4 |
| H21 | *Baptisia megacarpa* | Whitten 4647 | Bibb Co., AL | N/A | 469.5 |
| H32 | *Baptisia bracteata* | Bradley 3888 | Laurens Co., SC | N/A | 912.4 |
| H41 | *Thermopsis macrophylla* | N/A | MBOT - Living Collection -<br>2018-1231-2 | N/A | 3057.1 |
| H45 | *Thermopsis lupinoides* | N/A | MBOT - Living Collection -<br>2016-1242 | N/A | 6414.6 |
| H47 | *Thermopsis chinensis* | N/A | MBOT - Living Collection -<br>2020-0843 | N/A | 3571.4 |
| H48 | *Thermopsis robusta* | TEX - Mendenhall<br>682 | Siskiyou Co., CA | N/A | 1032.4 |
| H49 | *Thermopsis californica* | RSA-POM - White<br>10288 | Los Angeles Co., CA | N/A | 395.1 |
| NC2 | *Piptanthus nepalensis* | N/A | N/A | NC_056139.1 | N/A |
| NC6 | *Thermopsis lanceolata* | N/A | N/A | SRR8208358 | 4914.5 |
| NC10 | *Thermopsis turkestanica* | N/A | N/A | SRR12880891 | 5010.2 |
| NC11 | *Ammopiptanthus mongolicus* | N/A | N/A | SRR15039806 | 1343.3 |
| NC13 | *Piptanthus nepalensis* | N/A | N/A | SRR16038554 | 5092.3 |
| NC15 | *Euchresta tubulosa* | N/A | N/A | SRR14682996 | 912.4 |
| NC16 | *Lupinus albus* | N/A | N/A | SRR10902727 | 3057.1 |
| NC17 | *Crotalaria albida* | N/A | N/A | SRR18286790 | 6414.6 |
| RZ1 | *Thermopsis alpina* | N/A | N/A | PRJNA628236 | 1320.4 |
| ZH5 | *Piptanthus nepalensis* | N/A | N/A | SRR16038554 | N/A |

### DNA Isolation and Sequencing

Leaf materials were flash frozen in liquid nitrogen and then homogenized using a mortar and pestle. DNA isolations were conducted using the NucleoSpin Plant II Mini Kit for DNA from plants (Macherey-Nagel, Düren, Germany). Preliminary quality assessments were conducted using agarose gel visualization and NanoDrop (Thermo Fisher Scientific, Wilmington DE) quantification. When the minimum conditions for sequencing were met, samples were sent to the Beijing Genomics Institute (BGI; Shenzhen, China) where short-insert libraries (300 bp insert size) were prepared, and approximately 20 million paired end 150 bp reads with a Phred+33 encoded fastq quality score were generated using the MGI DNBseq platform (MGI Tech Co., Shenzhen, China). Adapter sequences, contamination, and low-quality reads were removed from the raw reads.

### Plastome Assembly

The complete plastome of one species and the raw read data of eight species were obtained from GenBank using SRA-toolkit v3.0.1, which were included among the 26 samples sequenced. *De Novo* draft plastome assemblies were prepared from each set of raw reads using the seed-and-extend method implemented in NovoPlasty v4.2, with the default *rbcL* elected as the starting seed (Dierckxsens et al., 2016). The raw reads of each taxon were mapped to their respective draft assemblies using Bowtie v1.3.0 (Langmead et al., 2009). Consensus assemblies were generated from the mapped sequences using the “generate consensus sequence” function in Geneious Prime v2021.1.1 (https://www.geneious.com). For each taxon, two alternative consensus sequences were generated, one with a 75% minimum per-site consensus threshold (no ambiguities), and the other with a 90% threshold (ambiguities present), resulting in two separate data sets.

### Plastome Alignment

Each of the two data sets were treated as follows: gene annotations were generated using the “live annotate & predict” function in Geneious Prime, with *Baptisia leucophaea* (Nutt.) Kartesz & Gandhi from Choi et al. (2022) used as a reference. Seventy-seven protein coding genes (CDS) were extracted from each plastome using the “extract annotation” function in Geneious Prime. Orthologous CDSs for each taxon were individually aligned using MAFFT v7.511 (Katoh and Standley 2013). Gaps were trimmed from each individual CDS alignment using TrimAL v1.2 (Capella-Gutierrez et al., 2009). All alignment columns containing ambiguities were masked using the “mask alignment” function in Geneious Prime. The individual trimmed and untrimmed alignments were concatenated, and NEXUS partition files were generated using catfasta2phyml.pl (Github: Nylander, 2010). Additionally, complete plastomes with one copy of the inverted repeat (IR) removed were aligned using MAFFT, and all gaps were trimmed using TrimAL, resulting in trimmed and untrimmed complete plastome alignments. A total of eight individual alignments were generated (Table 2).

**Table 2:** Summary of 6 nuclear (N1-N6) and 9 plastome (CP1-CP9) data sets utilized for phylogenomic inference. The lettered tree topologies correspond with those indicated in figure 3 with respect to the placement of *T. chinensis* and *V. turcica* relative to the North American clade.

| Dataset Name | Analysis Type | Data Type | Consensus Threshold (%) | Indels | Tree Topology |
| --- | --- | --- | --- | --- | --- |
| N1 | ML | 1540 Genes | 75 | + | A |
| N2 | ML | 1540 Genes | 75 | - | A |
| N3 | ML | 886 Genes | 75 | + | A |
| N4 | ML | 886 Genes | 75 | - | A |
| N5 | ASTRAL | 886 Genes | 75 | + | A |
| N6 | ASTRAL | 1540 Genes | 75 | + | A |
| CP1 | ML | Complete Plastomes | 75 | + | C |
| CP2 | ML | Complete Plastomes | 75 | - | C |
| CP3 | ML | Complete Plastomes | 90 | + | A |
| CP4 | ML | Complete Plastomes | 90 | - | A |
| CP5 | ML | 77 CDS Genes | 75 | + | B |
| CP6 | ML | 77 CDS Genes | 75 | - | B |
| CP7 | ML | 77 CDS Genes | 90 | + | B |
| CP8 | ML | 77 CDS Genes | 90 | - | B |
| CP9 | ASTRAL | 77 CDS Genes | 75 | + | A |

### Plastid Phylogenomics

Thirty-four taxa for which complementary nuclear data could be assembled were included in the analyses. For each alignment, a maximum likelihood (ML) analysis was conducted using IQtree v1.6.12 (Nguyen et al., 2015) with 1000 ultrafast bootstrap (BS) replicates (Hoang et al., 2018). Complete plastomes (minus one copy of IR) were treated as a single locus under the GTR+I+G substitution model. To accommodate rate heterogeneity in the CDS alignments, ModelFinder (Kalyaanamoorthy et al., 2017) was implemented. Additionally, individual gene trees for each CDS data set were generated using IQtree, and species trees consistent with multispecies coalescence were generated using ASTRAL-III (Zhang et al., 2018).

### Nuclear Gene Assembly

The individual sequences of five genistoid legume species were extracted from 1559 individual low-copy nuclear gene alignments generated by Zhao et al. (2021). The resulting sequences were used as references to generate low-copy nuclear genes from the raw read data of 33 taxa following a mapping-assembly-scaffold approach, implemented using the “assemb” and “ortho” options in PhyloHerb (Phylogenomic Analysis Pipeline for Herbarium Specimens; Cai et al., 2022). Raw reads were mapped to each reference gene using Bowtie v2.5.1, the resulting consensus sequences for each gene were assembled into contigs using Spades v3.15.5, and the resulting contigs were scaffolded using BLAST, with a default evalue of 1e-40. The individual nuclear gene sequences for the one reference species that corresponds with the plastome sequence obtained from GenBank (*Piptanthus nepalensis* Sweet) was added to the PhyloHerb output.

### Nuclear Gene Alignment

Each of the unaligned PhyloHerb output files were individually aligned using MAFFT v7.511. Two separate data sets were generated, one which included all 1540 gene alignments (more missing data), and another which included only the 886 genes that had complete taxon coverage (less missing data). Alternative data sets were generated from each of the resulting alignments – trimmed (all intragenic gaps removed) with TrimAl v1.2, and untrimmed (intragenic gaps present). The individual gene alignments of each of the resulting four data sets were concatenated using catfasta2phyml.pl (Table 2).

### Nuclear Gene Phylogenomics

Thirty-four taxa for which plastomes could be assembled were included in the analyses. For each nuclear data set, a ML analysis was conducted using IQtree v1.6.12 (Nguyen et al., 2015) with 1000 ultrafast bootstrap replicates (UFboot; Hoang et al., 2018). Due to the large size of each data set (34 taxa, each spanning ∼1.5 mbp), and the associated computational demands, the GTR+I+G substitution model was implemented, treating each concatenated data set as a single evolving locus. Additionally, individual gene trees for the intragenic-untrimmed data sets were generated using IQtree, and species trees consistent with multispecies coalescence were generated using ASTRAL-III (Zhang et al., 2018).

### Topology Testing

Two constrained trees were generated, each placing *T. chinensis* at the base of one of the two North American sub-clades. Using the -au option in IQtree, the topology of the constrained trees were tested against that of the unconstrained 886 nuclear gene trees (gaps present) with a bootstrap proportion test (bp-RELL), one sided Kishino-Hasegawa (KH) test, Shimodaira-Hasegawa (SH) test, expected likelihood weight (ELW) test, and an approximately unbiased (AU) test with 10,000 replicates (Kishino et al, 1990; Kishino and Hasegawa 1989; Shimodaira 2000; Strimmer and Rambaut 2002; Shimodaira, 2002). Additionally, the Siddal and Whitting method (SAW; 1999) was implemented by separately removing *V*. *turcica* and *T*. *chinensis* from the 886 nuclear gene alignment (Table 2: N3) and conducting a ML analysis on each of the resulting alignments.

### Divergence Time Estimation

Secondary calibrations were used to infer a time tree for the thermopsoids within a core-genistoids framework. To reduce the computational burden of Bayesian time tree inference, the nuclear loci data set was first subsampled to select for the most clock-like genes (Smith et al., 2018). Four hundred and twenty-seven nuclear gene trees were generated using IQtree, and then rooted by *Crotalaria albida* using the phyx program pxrr (Brown et al., 2017). The python script SortaData (Smith et al., 2018) was used to estimate the clock-likeness of each gene tree, measured according to bipartition scores, root-to-tip variance, and treelength. The top 15 genes were selected, and BEAST v2.7.1 was used to approximate divergence times based on a birth-death speciation prior and an optimized relaxed clock model of sequence evolution. The alignment data was partitioned by gene, with site models unlinked. The model averaging tool bModelTest (Bouckaert & Drummond, 2017) was implemented to select the most appropriate substitution model for each gene, and to measure the proportion of invariant sites and rate heterogeneity across sites. The standard substitution model package (SSM) v1.2.0 was used for model selection (Bouckaert & Xie, 2017). Three most recent common ancestor (MRCA) priors were derived from Lavin et al. (2005). Sigma values were adjusted to match the normal distribution of each prior to the respective mean and 95% highest density probability (HPD) of equivalent nodes in Lavin et al. (2005). The crown of the core genistoids (Crotalarieae – Genisteae + Sophoreae) was constrained to 45.5 Ma (95% HPD: 38.8 – 50.4 Ma), the crown of Crotalarieae – Genisteae was constrained to 41.2 Ma (95% HPD: 35.4 – 46.7), and the crown of the thermopsoid clade was constrained to 26.5 Ma (95% HPD: 18.4 – 34.5). Two independent Markov chain Monte Carlo (MCMC) chains were run, each for 1 × 10^6^ generations, with parameters logged every 1,000 generations. Tracer v1.7.2 was used to inspect MCMC convergence and effective sample size (>200) for each parameter. After discarding the first 10% of trees searched for each analysis as burn-in, trees were combined using LogCombiner v2.7.1, and the mean and 95% credibility intervals of most recent common ancestor nodes were calculated using TreeAnnotator v1.4.8.

### Biogeographic Inference

Ancestral area states were inferred based on a dispersal-extirpation-cladogenesis (DECLIKE) model of biogeography, implemented using the R package BioGeoBEARS v.1.1.2. (Matzke, 2013). The tips of the 15 nuclear gene tree generated in BEAST were scored according to three broad geographic localities: Eurasia (EE), Eastern North America (EN), and Western North America (WN). Analyses were conducted with and without the “+J” parameter. Each analysis utilized an areas adjacency matrix wherein North American localities were scored as 1 (adjacent) with respect to one another, and 0 (non-adjacent) with respect to Eurasia. To select which model conferred a higher likelihood on the data, likelihood ratio tests, as well as Akaike’s information criterion (AIC) and AICc (AIC corrected for small data sets) model weight tests, were conducted.

### Computation and Tree Visualization

Computational tasks were conducted using the Stampede2 supercomputer at the Texas Advanced Computing Center (TACC). Visualization and editing of phylogenomic trees were conducted using FigTree v1.4 (http://tree.bio.ed.ac.uk/software/figtree/) and Inkscape v1.2.1 (https://inkscape.org)

## Results

### Nuclear Phylogenomics

Nuclear-gene data sets were generated for a total of 34 taxa. From the original 1559 reference genes, the per taxon number of genes recovered ranged from 886 to 1540, and the per taxon concatenated gene sequence length ranged from 838 kb to 1.281 mb. Though the sequence alignments prepared for ML analysis varied considerably in the number of parsimony informative sites (PIS), ranging from 14,148 (Table 2: N2) to 164,236 (Table 2: N1), the resulting backbone topologies and corresponding bootstrap support values were congruent across all nuclear data sets. *Ammopiptanthus* formed a clade with *Euchresta*, placing this genus outside of the core thermopsoids (Fig. 1, Group E). Among the core thermopsoids, a fully resolved clade composed exclusively of Eurasian taxa was supported (Fig. 1, Group D). Within this clade, *Piptanthus* formed a subclade with *T. lanceolata* R. Br, *T. turkestanica* Gand, and *T. dahurica* Czefr, sister to *T. alpina* Ledeb. *Vuralia turcica* formed a clade with *T. chinensis*, sister to the North American thermopsoids (Fig 1; Group C). The North American taxa formed a strongly supported clade composed of two subclades, one comprising all members of *Baptisia* (Fig 1; Group B), and the other comprising the North American *Thermopsis*, and the Asian *T. lupinoides* (Fig 1; Group A). The results of the nuclear-based multispecies coalescence analyses were strongly congruent with the nuclear-based ML analyses, except for the placement of *Ammopiptanthus*, which was sister to the core thermopsoid clade, making its inclusion within this group ambiguous (Fig. 2).

**Figure 1:**
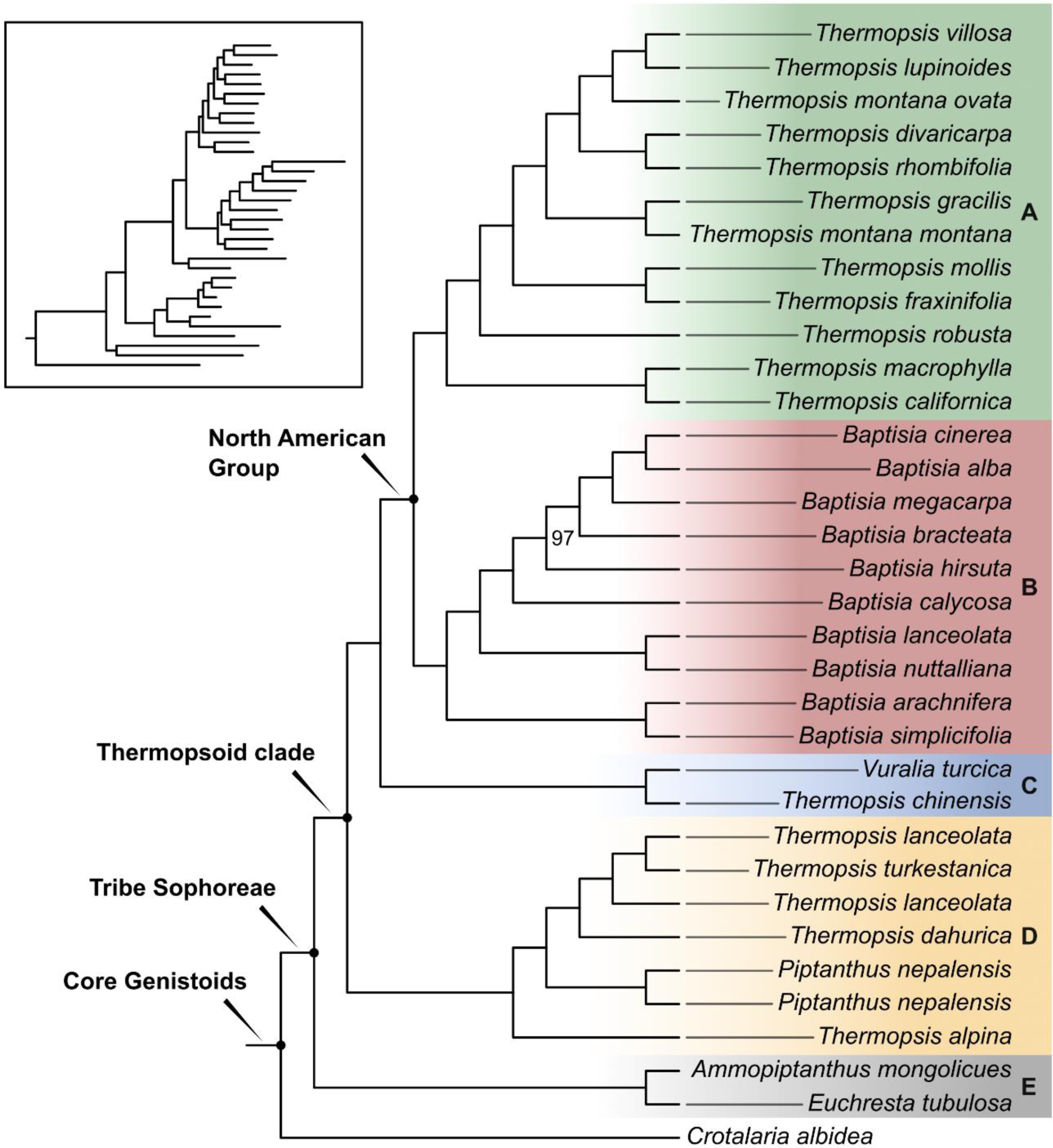
Maximum likelihood cladogram of 886 nuclear genes with complete taxon coverage (N3, Table 2). Boot strap support values less than 100 are indicated. Branch lengths are indicated in the untransformed tree shown in the upper left corner. Each major clade is indicated by letters, including the North American *Thermopsis* clade (A), the *Baptisia* clade (B), the *T. chinensis – V. turcica* clade (C), the Eurasian thermopsoid clade (D), and the sophoroid clade (E).

**Figure 2:**
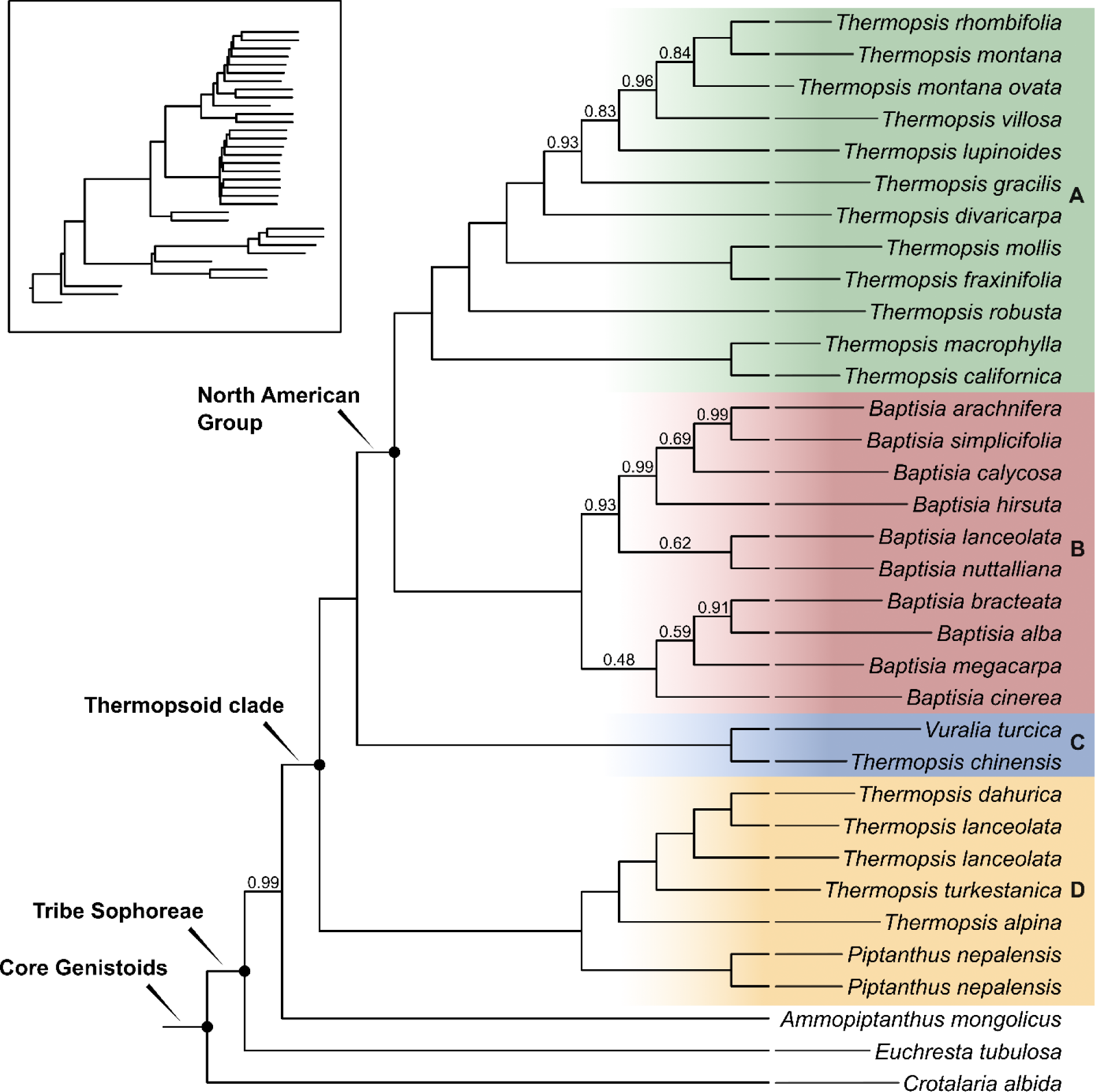
ASTRAL species tree based on 1540 nuclear gene trees (N6, Table 2). Local posterior probability values less than one are indicated. Branch lengths are indicated in the untransformed tree shown in the upper left corner. Each major clade is indicated by letters, including the North American *Thermopsis* clade (A), the *Baptisia* clade (B), the *T. chinensis – V. turcica* clade (C), and the Eurasian thermopsoid clade (D).

### Topology Testing

Given the unexpected placement of *T. chinensis* as sister to *V. turcica*, outside of the North American clade where it is traditionally placed, two alternative topologies were tested, each placing *T. chinensis* at the base of one of the two North American clades, as seen in the ML analyses of Zhang et al. (2015) and Shi et al. (2017). The KH, SH and AU tests each returned p-values of < 0.001, significantly excluding *T. chinensis* from placement within the North American clade. This finding was supported by the results of the log-likelihood difference, bootstrap proportion, and expected likelihood weight tests (Table S1). Furthermore, to explore the potential effects of long-branch attraction (LBA) on the placement of *T. chinensis*, separate modifications were made to the N3 (Table 2) data set, one in which *V. turcica* was excluded and the other in which *T. chinensis* was excluded. The reciprocal exclusion of either taxon did not affect the relative placement of the other, thus rejecting the LBA hypothesis (Figure S2).

### Plastome Phylogenomics

Thirty-three plastomes were assembled and included with a previously assembled plastome obtained from GenBank (*Piptanthus nepalensis*). Plastome length varied from 153,767 bases (*T. lanceolata*) to 150,757 bases (*V. turcica*). Depth of coverage ranged from 316 to 6,414, with an average of 2,246 (Table 1). All ingroup plastomes were colinear. Relative to the ingroup, *Crotalaria albida* was found to have a ∼23 kb inversion in the large single copy region. The PIS values of sequence alignments ranged from 447 (Table 2: CP5) to 5950 (Table 2: CP1). Unlike the nuclear-based ML analyses, the tree topologies inferred by the plastid-based ML analyses varied considerably between data sets, particularly with respect to the placement of the *T. chinensis – V. turcica* clade. Data sets CP5 – CP8 (Table 2) each produced a tree topology reflected by Fig. 3B, in which the *T. chinensis – V. turcica* clade was within the North American clade, sister to *Baptisia*. Data sets CP1 – CP2 each produced a tree topology reflected by Fig. 3C, in which the *T. chinensis – V. turcica* clade was sister to the North American *Thermopsis*. When the read-mapping consensus threshold was increased to 90% (CP3 – CP4), from the standard 75%, a backbone topology congruent with that of the nuclear-based analyses was produced (Fig. 3A). The plastome-based ASTRAL analysis (CP9) also place *V. Turcica* and *T. chinensis* outside of the North American clade (Figure S15).

**Figure 3:**
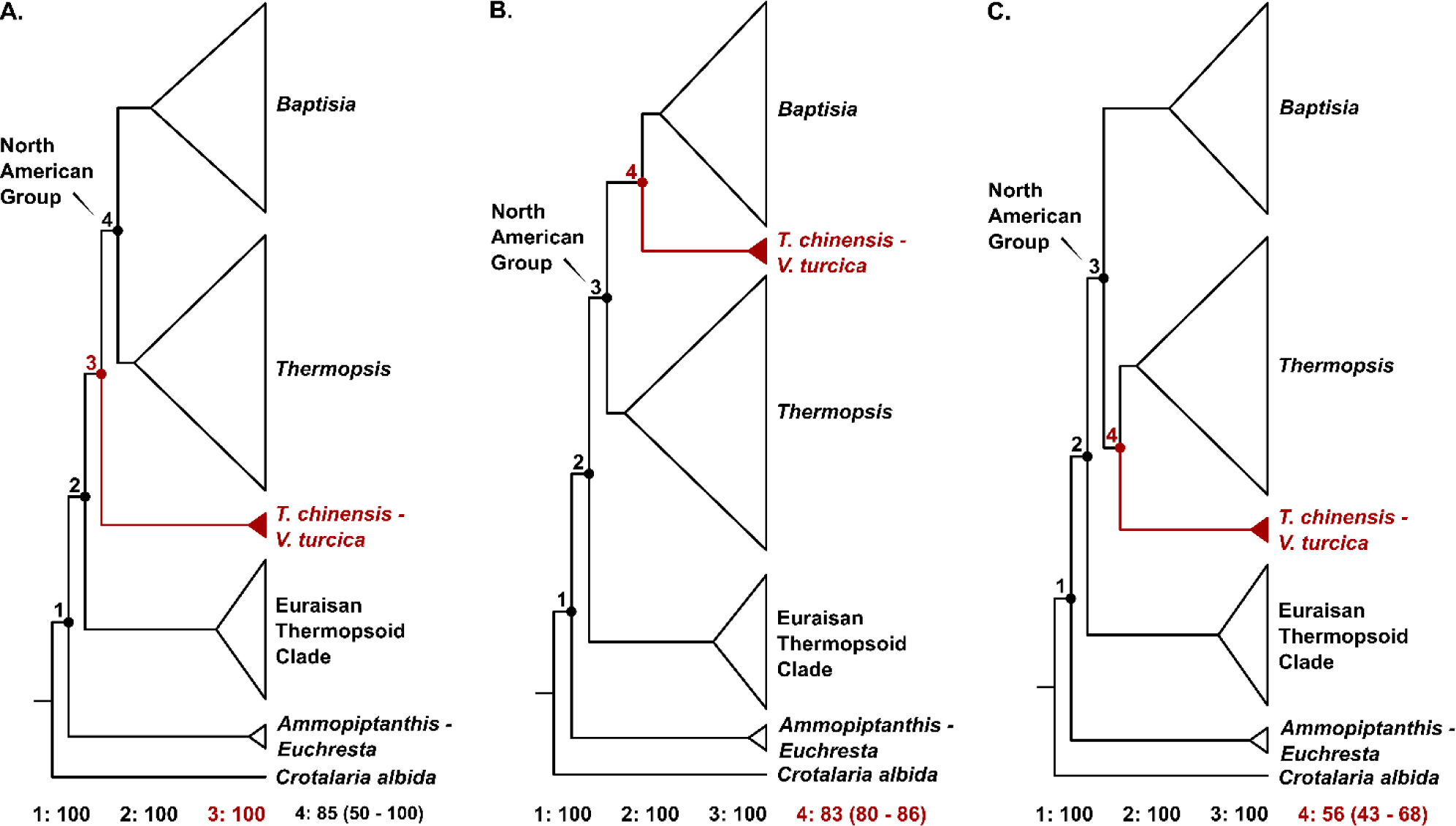
Three tree topologies that differ in the placement of the *T. chinensis* – *V. turcica* clade generated from 12 ML analyses, based on either 886 nuclear genes, 1540 nuclear genes, 77 plastome CDS genes, or complete plastomes. Tree A is supported by six analyses, including all nuclear-based analyses and a subset of plastome-based analyses, trees B is supported by four plastome-based analyses, and tree C is supported by two-plastome based analyses. The average bootstrap values for all analyses that support each numbered node of respective tree are indicated below each tree, with the ranges of boot strap support values indicated in parentheses when less than 100. The specific analyses that correspond with each tree topology are summarized in Table 2 (N1 – N4 and CP1 – CP8).

### Divergence Time Estimation

To explore the biogeographic implications of the *T. chinensis – V. turcica* clade, divergence time estimation and ancestral area state reconstruction were conducted using the N3 data set. A representative of the Genisteae clade, *Lupinus albus*, was added to the data set to accommodate a total of three secondary calibration points. The addition of *L. albus* to the nuclear data reduced the number of genes with complete taxon coverage from 886 to 427; clock-likeness testing further reduced this number to 15. Maximum likelihood analysis of the resulting 15 gene data set placed *Lupinus* in a clade with *Crotalaria*, thus informing the topology at the constrained calibration nodes of the BEAST analysis. The *T. chinensis* – *V. turcica* clade was fully supported as sister to the North American clade. The results of the BEAST analysis (Fig. 4) indicated a crown core genistoids at 46.4 Ma (95% HPD: 40.8 – 52.2 Ma), and crown Sophoreae at 29.9 Ma (95% HPD: 22.9 – 37.8 Ma). The thermopsoid clade was shown to have emerged at 23.9 Ma (95% HPD: 18.0 – 29.7 Ma), diverging into the Eurasian thermopsoid clade, with a crown age of 11.1 Ma (95% HPD: 6.7 – 16.4 Ma), and the *T. chinensis – V. turcica* + North American clade, with a crown age of 12.3 Ma (95% HPD: 7.9 – 17.0 Ma). The north American lineage was shown to have diverged into the North American *Thermopsis* and *Baptisia* at 8.9 Ma (95% HPD: 5.7 – 12.38 Ma).

**Figure 4:**
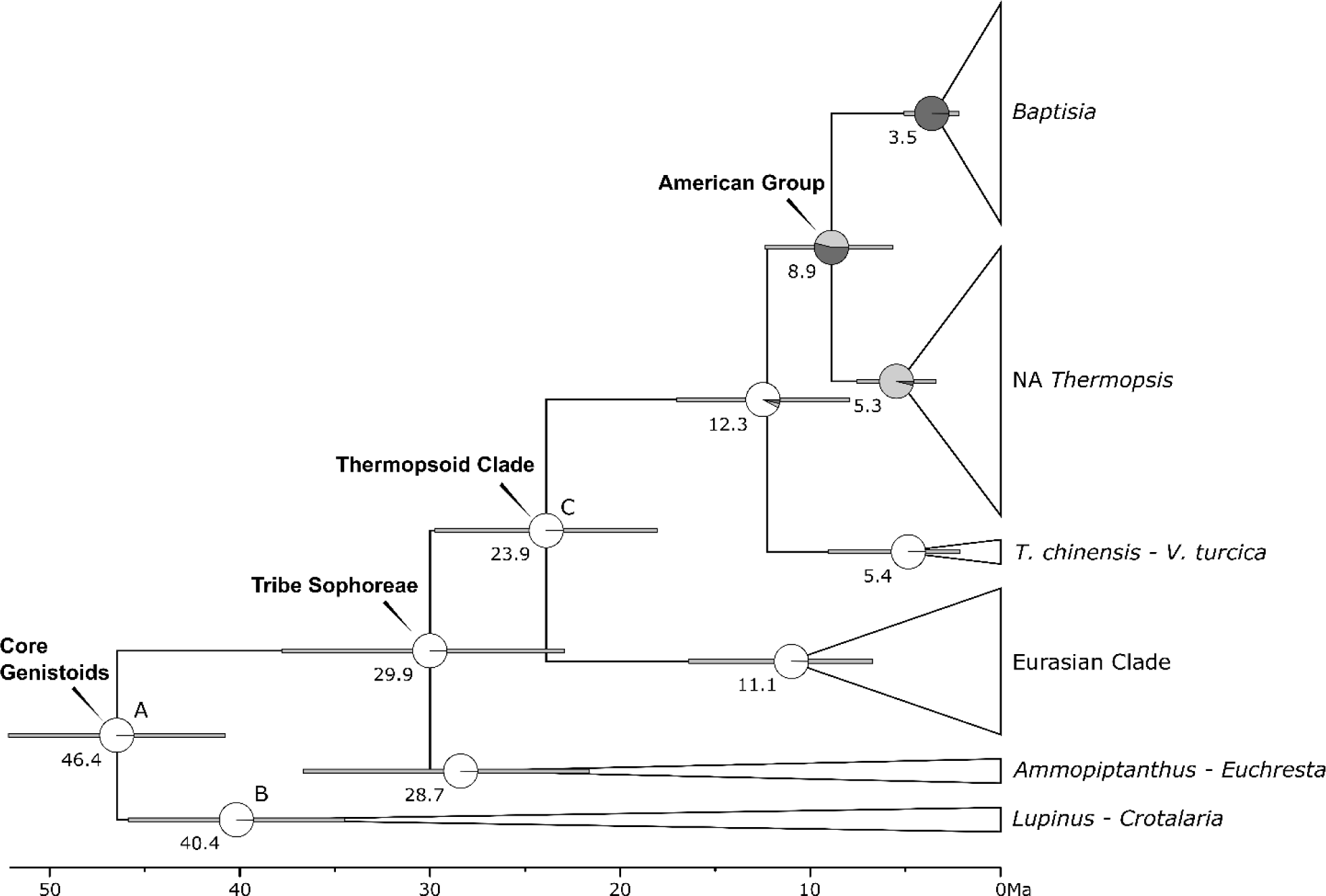
Bayesian BEAST chronogram based on 15 clock-like nuclear genes. Divergence times in millions of years (Ma) are indicated at nodes. Four secondary calibration points are indicated by letters: A (45.5 Ma), B (41.2 Ma), and C (26.5 Ma). Ninety-five percent highest density probability (HPD) values are indicated by gray bars at nodes. Ancestral area probabilities, based on the DECLIKE+J criterion, are indicated by pie charts at each node, with white indicating a Eurasia origin, medium gray indicating a western North America origin, and dark gray indicating an eastern North America origin. Posterior probabilities were unanimous across the tree.

### Biogeographic Reconstruction

Likelihood ratio tests strongly supported the alternate hypothesis that the addition of the +J parameter (p < 0.00001), which allowed for founder/jump speciation events, conferred a higher likelihood on the data (Table S2; Matzke 2013). All Old World thermopsoid lineages, including the ancestor to the *T. chinensis – V. turcica* clade + North American clade, were shown to be of Eurasian origin. *Baptisia* and the North American *Thermopsis*, as well as the common ancestor between them, were shown to be of North American origin. The ancestor of *T. lupinoides*, the single old-world taxon nested within the North American clade, was also shown to be of North American origin. Though the ancestor of the North American clade was shown to be of mixed Eastern/Western North American origin, a slight Eastern North America bias was detected (Fig. 4).

## Discussion

### Thermopsideae Phylogenomics

The ASTRAL and ML analyses each placed *Ammopiptanthus* sister to the thermopsoid clade, with the ML analyses placing the genus in a fully supported clade with *Euchresta*, well outside of the traditional Thermopsideae. These results support the conclusions of Cardoso et al. (2013), which merged Thermopsideae with Sophoreae *sensu* Cardoso, making the group a monophyletic member of the core genistoids. Though a more inclusive Sophoreae avoids taxonomic over-fragmentation, the names sophoroid and thermopsoid are adopted from Shi et al., (2017) to describe two well supported Sophoreae clades identified by the present study. All nuclear-based and a subset of plastome-based analyses support three thermopsoid clades – a North American clade, a Eurasian clade, and a *T. chinensis – V. turcica* clade. The thermopsoids (excluding *Pickeringia*) comprise *Anagyris*, *Baptisia*, *Piptanthus*, *Thermopsis*, and *Vuralia*. The genera of the sophoroid clade included in the present study are *Ammopiptanthus* and *Euchresta* (Fig. 1).

### Eurasian Thermopsoid Clade

The Eurasian thermopsoid clade (Fig 1: Group D) is composed of *Thermopsis, Piptanthus, and Anagyris*. Despite the cytonuclear discordance observed within the group, both nuclear and plastome based analyses supported a subclade comprising the widespread Eurasian *T. lanceolata*, the Mongolian *T. dahurica*, and the Central Asian *T. turkestanica.* Consistent with the findings of Shi et al., (2017), the present ML analyses nested *Piptanthus* within the Eurasian clade, with the nuclear-based analyses placing the genus in a clade with the *T. lanceolata* – *T. turkestanica* group, sister to *T. alpina,* and the plastome-based analyses placing the genus in a clade with *T. alpina*, sister to the *T. lanceolata* – *T. turkestanica* group. Both nuclear based ASTRAL analyses (Fig. 2: Group D; Fig. S6) indicated a monophyletic *Piptanthus* as sister to the rest of the Eurasian taxa, whereas the plastome based ASTRAL analysis nested the genus with *T. alpina*, consistent with the plastome based ML analyses (Fig. S15). Though the Mediterranean *Anagyris* was not included in these analyses, past phylogenetic studies have placed this genus in the Eurasian clade, closely allied with the *T. lanceolata* – *T. turkestanica* group (Wang et al. 2006; Shi et al., 2017). These results did not support the monophyly of *Thermopsis*.

### T. chinensis – V. turcica Clade

A strongly supported clade comprising the Turkish endemic *V. turcica* and the east Asian endemic *T. chinensis* was sister to the North American group. An association between *T. chinensis* and the North American *Thermopsis* was first noted by Czefranova (1970), which focused on morphological series within *Thermopsis*. This finding was later contradicted by the quinolizidine alkaloid studies of Salatino and Gottlieb (1980), resulting in the exclusion of *T. chinensis* from early cladistic studies of the North American Thermopsideae (Mendenhall 1994). Nevertheless, subsequent phylogenetic studies placed this taxon firmly within the North American clade, suggesting it to be the result of long-distance North America–East Asia dispersal. This interpretation, however, was not supported by the present study, as all nuclear-based analyses, and a subset of plastome-based analyses supported a *T. chinensis* – *V. turcica* clade, positioned outside of the North American clade.

*Vuralia turcica*, an endemic of marshy areas surrounding the Eber lakes in Aksehir, Turkey, was first described in 1983 under the name *Thermopsis turcica* Kit Tan, Vural & Küçük, but later recircumscribed into a monotypic *Vuralia* by Uysal et al. (2014), based on morphological autapomorphies such as a three-carpellate ovary and indehiscent fruit, as well as an ITS analysis which placed the genus distant from the other thermopsoids. Uysal et al. (2014) did not include *T. chinensis* in their study, however the subsequent ITS analysis of Shi et al. (2017), that included both *V. turcica* and *T. chinensis*, showed the two taxa to form a clade, nested within the North American group. Though the present results placed the *T. chinensis* – *V. turcica* clade outside of the North America group, contradicting the ML studies of Shi et al. (2017), neither study supported the monotypic generic status of *Vuralia*.

The contradictory placement of *T. chinensis* – *V. turcica* likely stems from the relatively short internal branch between the *T. chinensis* – *V. turcica* clade and North American clade. This feature, when combined with the limited number of genetic markers analyzed by previous studies, may have resulted in either a collapsed internal branch, or the long branch attraction of *T. chinensis* to the base of either of the North American subclades (Shi et al., 2017). In the present study, the *T. chinensis* – *V. turcica* clade is fully resolved and strongly supported across all nuclear ML analyses, and the LBA hypothesis is rejected. Though the subset of analyses based on a 75% read mapping consensus threshold for complete plastomes placed the *T. chinensis* – *V. turcica* clade within the North American group, an increase to a consensus value of 90% resulted in trees congruent with the nuclear analyses, suggesting this finding to have resulted from the signal obscuring effects of plastid heteroplasmy (Gonçalves et al., 2020).

The small size of the *T. chinensis – V. turcica* clade, compared to its much larger North American counterpart, as well as the relatively short internal branch connecting the two groups, suggests the *T. chinensis* – *V. turcica* clade to be a relic of the Asian ancestor that gave rise to the North American group, a finding that raises questions regarding the degree to which intercontinental genetic exchange may have occurred during the early divergence between the East Asian and North American lineages.

### North American Clade

In his 2006 treatment of *Baptisia*, Turner identified five morphological series within the group, informed largely by the morphometric work of Mendenhall (1994). To test these groupings, two individuals of each series were included in the present analyses, along with a single individual for all North American species of *Thermopsis*, informed by the taxonomic treatment of Chen et al. (1994). A relatively high degree of cytonuclear discordance was detected between nuclear and plastome data sets (Fig. S1). Despite this, all analyses supported a monophyletic North American thermopsoid group, divided into two strongly supported subclades, one comprising *Baptisia*, and the other comprising the North American *Thermopsis*, as well as a single East Asian taxon, *T. lupinoides*. Within each subclade, BS support values resulting from plastome based analyses were considerably lower than those of the nuclear based analyses, likely due to a relatively lower number of parsimony informative sites and shorter average pairwise genetic distance in the plastome data sets. Additionally, a high degree of interspecific hybridization and regional intergradation has been reported among *Baptisia* and the North American *Thermopsis*, potentially contributing to the high level of observed discordance (Alston and Turner, 1963; Leebens-Mack and Milligan, 1998). Given these challenges, caution was taken when interpreting intrageneric relationships in the North American group, and the nuclear based relationships with the greatest overall support (Fig. 1) are emphasized.

Across all data sets, only a single subclade of *Baptisia*, comprising *B. lanceolata* Elliott and *B. nuttalliana* Small, supported the morphological groupings of Turner (2006). Excluding the results of plastome-based analyses, an additional group, the “simple leaf” clade, comprising *B. arachnifera* W. H. Duncan and *B. simplicifolia* Croom, was strongly supported by both the ASTRAL (Fig. 2) and ML analyses (Fig 1), with the latter analysis showing this group to be the earliest diverging lineage in *Baptisia*. Four of the remaining six taxa were placed in a single subclade by both nuclear and plastome based ML analyses, including *B. hirsuta* Small, and *B. megacarpa* Chapm. ex Torr. & A. Gray. *Baptisia bracteata* Muhl. Ex Elliott and *B. cinerea* (Raf.) Fernald & B. G. Schub, which formed a third morphological group in Turner (2006), were also included in the subclade. The remaining two members of *Baptisia*, *B. calycosa* Engelm and *B. alba* R. Br., were highly discordant across all analyses, making their phylogenomic positions within the group unclear.

Within the North American *Thermopsis* clade, a single subclade comprising *T. macrophylla* Hook. & Arn. and *T. californica* S. Watson was supported by both nuclear and plastome based ML (Fig. 1A) and ASTRAL analyses (Fig. 2), which indicated this group to be the earliest diverging North American *Thermopsis* lineage. A *T. californica – T. macrophylla* grouping is not surprising given that the two taxa have been considered synonyms of *T. macrophylla* by previous works, including the treatment of Isely (1981), which considered the two, along with *T. robusta* Howell, and *T. gracilis* Howell, to be regional variants. The nuclear based ML and ASTRAL analyses strongly supported a second clade, comprising *T. mollis* (Michx.) M. A. Curtis and *T. fraxinifolia* M. A. Curtis, two of the three eastern North American members of *Thermopsis*. This grouping was noted by Czefranova (1970) who combined the two taxa to form series Fraxinifoliae. Others, including Isely (1981), have considered the two to be synonyms of *T. fraxinifolia*. The third eastern *Thermopsis*, *T. villosa* (Walter) Fernald & B. G. Schub, is shown to be distant from the *T. mollis* – *T. fraxinifolia* clade, suggesting two independent intracontinental vicariance events involving the North American *Thermopsis*. The remaining members of *Thermopsis* are highly discordant across all analyses, making their phylogenomic positions ambiguous. It is worth noting that the nuclear and plastome based ML and ASTRAL analyses nested a single Asian taxon, *T. lupinoides*, among the North American *Thermopsis*, sister to either the easternmost or westernmost members, *T. villosa*, or *T. gracilis*, respectively. This finding supports the previously reported North America – East Asia disjunct status of the North American *Thermopsis*, a biogeographic pattern which is rare among Papilionoideae legumes (Sa et al., 2000; Wang et al., 2006; Uysal et al., 2014; Zhang et al., 2015; Shi et al., 2017).

### Thermopsoid Biogeography

Uplift of the Qinghai-Tibetan Plateau (QTP), located at the intersection of Central, South, and East Asia, has been implicated by previous studies as the principal driver of early diversification in Thermopsideae (Zhang et al., 2015, Shi et al., 2017). Prior works have identified three major phases of QTP uplift, resulting from continental collision along the convergent boundary between the Indo-Australian and Eurasian tectonic plates (Wang et al., 2018). The first phase (35 – 55 Ma) gave rise to the coniferous Gangdise mountains, the second phase (30 – 20 Ma), involving the westward withdrawal of the Paratethys Sea, resulted in the aridification of the Asian interior, and a third phase (15 – 8 Ma), involving the rapid uplift of the plateau margins, resulted in the development of the East Asia monsoons (Zhisheng et al., 2001; Dupont-Nivet et al., 2009; Wang et al., 2018).

A number of studies concerning the age and biogeographic distribution of the thermopsoids have been conducted, including Lavin et al. (2005), which dated the mean crown age of the thermopsoid clade at 26.5 Ma, Zhang et al. (2015), which indicated a central Asia origin of the thermopsoid clade, dated at 20.32 Ma, and Shi et al. (2017), which dated the group at 24.5 Ma. The present nuclear-based ancestral area reconstruction and dating analyses strongly support a Eurasian origin of the thermopsoid clade, with a stem age of 29.9 Ma, and a crown age of 23.9 Ma (Fig. 4). The relative disparity between the present results, and those of Zhang et el. (2015), likely arise from the phylogenomic placement of *Ammopiptanthus*, with the former placing the genus within the sophoroid group, and the latter placing the genus in a clade with the thermopsoids, distant from *Sophora*. Nevertheless, the present results support the hypothesis put forth by Zhang et al. (2015), namely that the second phase of QTP uplift, which drove the central Asian climate toward continental conditions, likely influenced early diversification of the thermopsoids, driving dispersals into the QTP, central Asia, and across the Bering Strait into North America.

Two equally parsimonious biogeographic hypotheses were inferred from the placement of the *T. chinensis – V. turcica* clade sister to the North American clade: the ancestor of the *T. chinensis – V. turcica* clade and the North American clade arose in North America, followed by two long distance dispersal events; one involved dispersal into Asia by the ancestor of the *T. chinensis – V. turcica* clade, and the other involved dispersal into Asia by the ancestor of *T. lupinoides*. Alternatively, the ancestor of the *T. chinensis – V. turcica* clade and the North American clade arose in Asia, followed by two-long distance dispersal events. Of these events, one involved dispersal into North America by the ancestor of the North American clade, and the other involved a subsequent dispersal into Asia by the ancestor of *T. lupinoides*. The present analyses support the latter hypothesis, indicating a Eurasian origin of the ancestor between the *T. chinensis – V. turcica* clade and the North American clade. This finding is shown to greatly impact the stem to crown age of the North American clade. For instance, in Zhang et al. (2015), the North American lineage, with a crown age of 5.48 Ma, diverged from its most recent Asian ancestor ca. 20.32 Ma. However according to the present study, the North American clade, with a mean crown age of 8.9 Ma, diverged from its most recent Asian ancestor ca. 12.3 Ma, reducing the timing of East Asia–North America dispersal to a 3.4-million-year window in the late Miocene, corresponding with an increase in aridity of the Asian interior, as well as the onset of Indian and East Asian monsoons (Fig. 4; Zhisheng 2001). Such East Asia–East North America floristic exchanges are commonly reported, presumably due to similarities in the ecology between the two regions (Liu et al., 2017). In a number of cases, the estimated timing of dispersal aligns with the present findings, such as *Cornus* 13.1 Ma (Xiang et al., 1998), *Phaseolus* 12.5 Ma (Jabbour et al., 2018), and *Epimedium-Vancouveria* 9.7 Ma (Zhang et al., 2007). Furthermore, it is often the case that East Asia–East North America disjunct groups exhibit a greater degree of biodiversity among their Asian lineage, as exemplified by the genus *Gleditsia*, for which there are eight East Asian species, and two North American species (Schnabel et al., 2003). Perplexingly, the opposite is true for the North America thermopsoid clade + *T. chinensis – V. turicica* clade, where the North American lineage radiated into ca. 27 species, while its sister Asian lineage appears to have been driven to near extinction.

### Taxonomic Implications

The present study indicated a polyphyletic *Thermopsis*, as noted by numerous previous works, highlighting the need for taxonomic revision within the group (Wang et al. 2006; Zhang et al., 2015; Shi et al., 2017). The merging of the thermopsoid genera into an inclusive *Thermopsis* would make the group monophyletic and avoid the issue of taxonomic over-fragmentation. However, given the degree of morphological and geographic divergence between the major thermopsoid lineages, this interpretation would be of little taxonomic value. Furthermore, the generic concept of *Thermopsis*, typified by *T. lanceolata* R.Br. (1828), is predated by the genus *Anagyris* L. (1753), which has been shown by the present study to form a well-supported clade with *T. lanceolata*. Consequently, the merging of *T. chinensis* with *Vuralia*, and the rest of the Eurasian thermopsoids with *Anagyris*, would split the old-world taxa into two monophyletic clades, while satisfying the traditional rules of botanic nomenclature. In this view, the North American *Thermopsis* would be transferred to the synonym *Thermia* (Nuttall 1818), and *Baptisia* would retain its generic status. Additional phylogenomic work, including a complete sampling of the old world thermopsoids, as well as a detailed analysis of morphology both within and between the major thermopsoid lineages, is needed in order to better inform future taxonomic revision within the group.

## Conclusion

The thermopsoids exhibit a number of highly unusual biogeographic features, including an East Asia– West Asia disjunction, two independent and bidirectional East Asia–North America disjunctions, as well as at least two independent Western North America–Eastern North America disjunctions. These features, combined with the relatively high degree of cytonuclear discordance observed within the North American lineages, as well as the high degree of observed polyphyly in *Thermopsis*, raise the need for more rigorous genome-scale testing. Additionally, future studies seeking to clarify the taxonomy of the group and improve resolution of the North American thermopsoid clade will benefit from more complete taxon sampling, which in turn should support a better understanding of factors that drove the numerous long-distance dispersals within the group.

## Acknowledgements

This study was financially supported by the Texas Ecological Laboratory Program and the Department of Integrative Biology at the University of Texas at Austin. Thanks is given to Dr. Liming Cai, Dr. George Yatskievych, and Dr. Tracey Ruhlman for their comments and feedback regarding earlier versions of this manuscript, and to Dr. Liming Cai, Dr. Edward Theriot, Dr. David Cannatella, and Dr. Domingos Cardoso for suggesting analyses. Thanks is also given to the Billie L. Turner Plant Resources Center at the University of Texas at Austin, the Missouri Botanic Garden, and the California Botanic Garden, for supplying leaf materials for taxa critical to this study.

