## Supplementary material for "Out of Place: Three Eurasian Taxa Shed Light on Origin of Thermopsideae in North America"

**Supplementary Tables and Figures:**

**Table S1.** Results of ML tree topology testing based 886 nuclear genes, conducted in IQtree. *Thermopsis chinensis* was constrained to either the base of the *Baptisia* clade, or to the base of the North American *Thermopsis* clade. Each test statistically rejected the constrained topologies and supported the unconstrained topology, wherein *T. chinensis* is placed in a clade with *V. turcica*, sister to the North American group.

| Tree | LogL | deltaL | bo(-)RELL | p(-)KH | p(-)SH | c(-)ELW | pAU |
|---|---|---|---|---|---|---|---|
| Unconstrained | -4654634.72 | 0 | 1 (+) | 1 (+) | 1 (+) | 1 (+) | 1 (+) |
| Sister to Baptisia | -4661716.78 | 7082.1 | 0 (-) | 0 (-) | 0 (-) | 0 (-) | 1.95E-09 (-) |
| Sister to NA Thermopsis | -4661697.39 | 7062.7 | 0 (-) | 0 (-) | 0 (-) | 0 (-) | 3.03E-13 (-) |

deltaL : logL difference from the maximal logl in the set.
bp-RELL : bootstrap proportion using RELL method (Kishino et al. 1990).
p-KH : p-value of one sided Kishino-Hasegawa test (1989).
p-SH : p-value of Shimodaira-Hasegawa test (2000).
c-ELW : Expected Likelihood Weight (Strimmer & Rambaut 2002).
p-AU : p-value of approximately unbiased (AU) test (Shimodaira, 2002).

Plus signs denote the 95% confidence sets.
Minus signs denote significant exclusion.
All tests performed 1000 resamplings using the RELL method.

**Table S2.** Analyses of ancestral area states were conducted in BioGeoBears under three different statistical criteria, each with (null) and without (alt) the +J parameter. In all three cases p values (pval) of < 0.00001 were returned, thus supporting the alternate hypothesis that the addition of the +J parameter confers a higher likelihood on the data.

| null | alt | LnLalt | LnLnull | DFalt | DFnull | DF | Dstatistic | pval |
|---|---|---|---|---|---|---|---|---|
| DEC | DEC +J | -20.21 | -39.9 | 3 | 2 | 1 | 39.37 | 3.50E-10 |
| DIVALIKE | DIVALIKE +J | -20.23 | -38.47 | 3 | 2 | 1 | 36.46 | 1.60E-09 |
| BAYAREALIKE | BAYAREALIKE +J | -20.23 | -53.88 | 3 | 2 | 1 | 67.29 | 2.30E-16 |

**Figure S1:** Mirrored ML cladograms of 886 nuclear genes (N3; Left, Table 2) and complete plastomes (CP3; Right, Table 2). BS support values where < 100 are shown at nodes. Phylograms that correspond with each cladogram are shown in the upper corners.

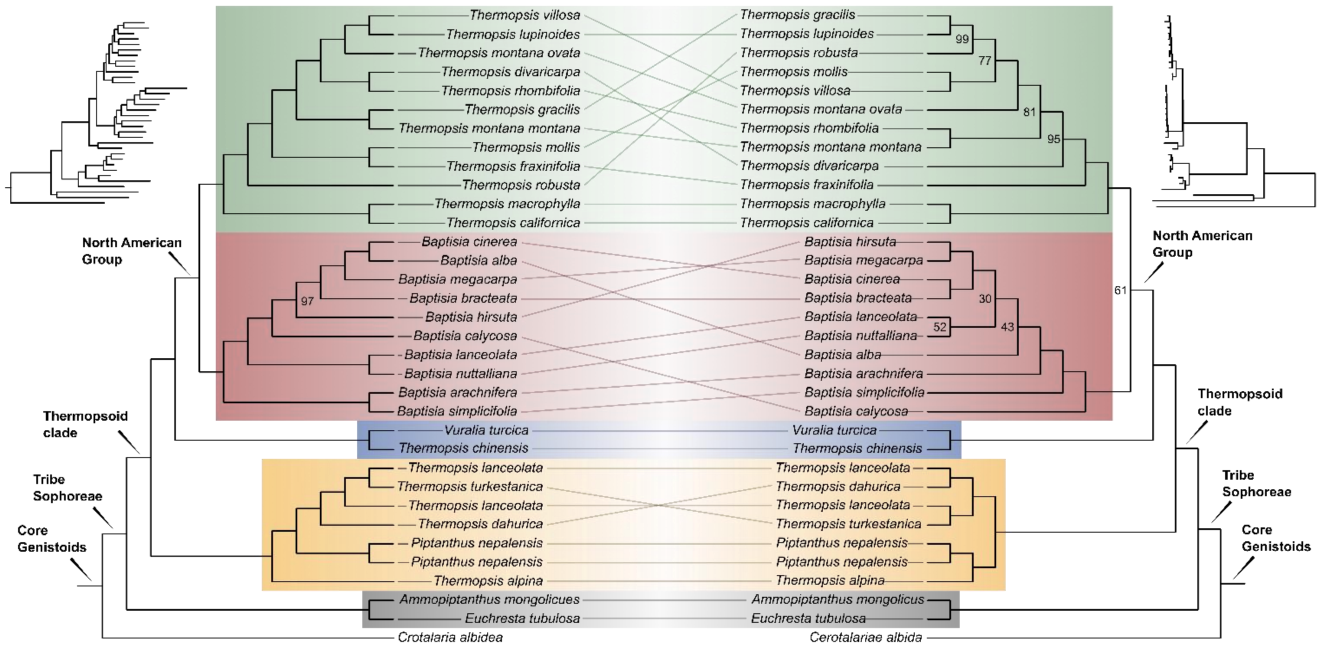

**Figure S2:** The LBA hypothesis was tested by removing *T. turcica* from the N3 data set (Table 2) and conducting a maximum likelihood analysis using IQtree. Bootstrap support values based on 1000 ultrafast bootstrap replicates are indicated at each node. The removal of *V. turcica* was not shown to affect the placement of *T. chinensis* (H47) outside of the North American clade. Codes for taxon names are in Table 1.

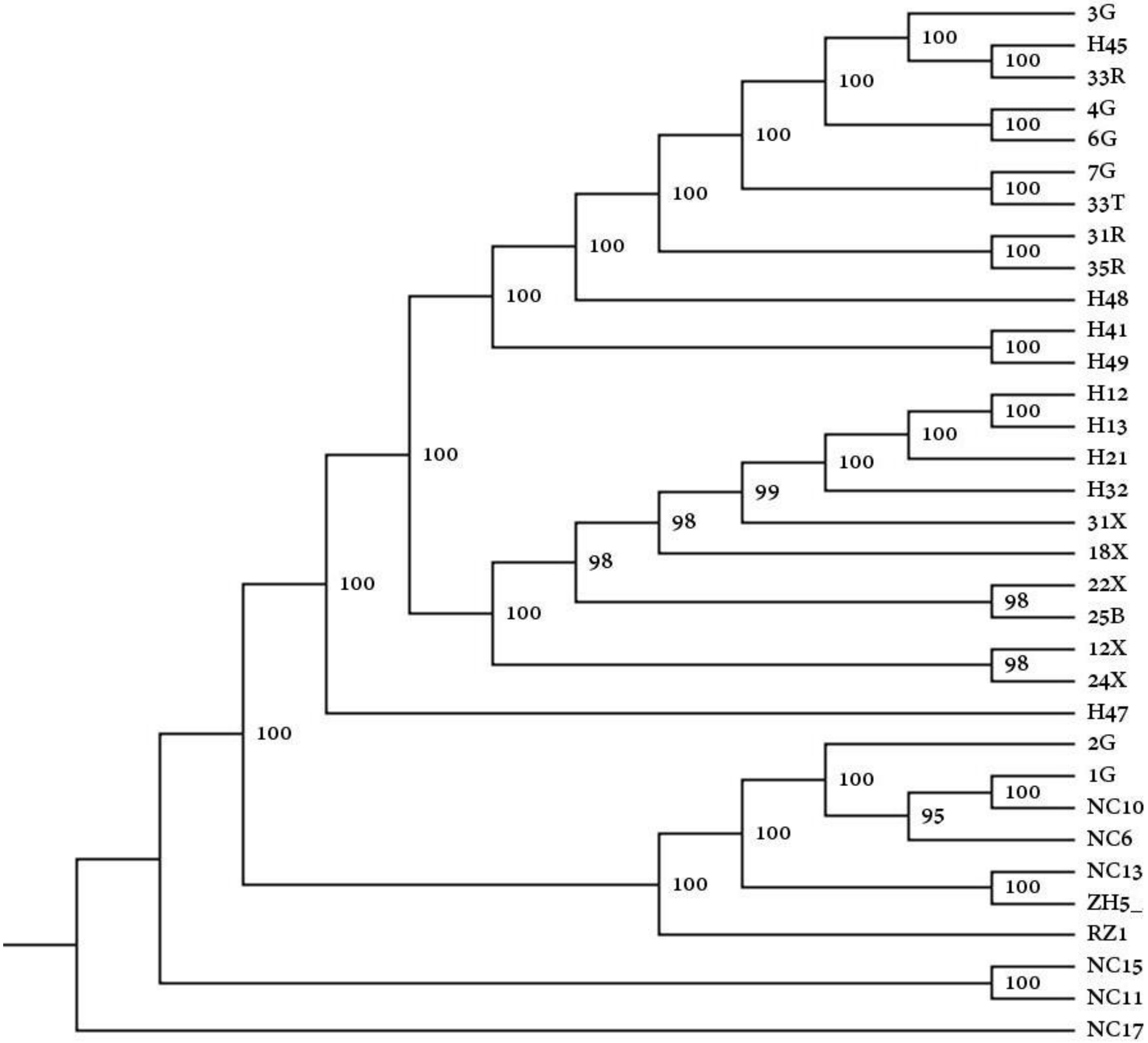

**Figure S3:** Maximum Likelihood cladogram based on 1540 nuclear genes corresponding with the N1 data set (Table 2), generated using IQtree. Bootstrap support values based on 1000 ultrafast bootstrap replicates are indicated at each node. Codes for taxon names are in Table 1.

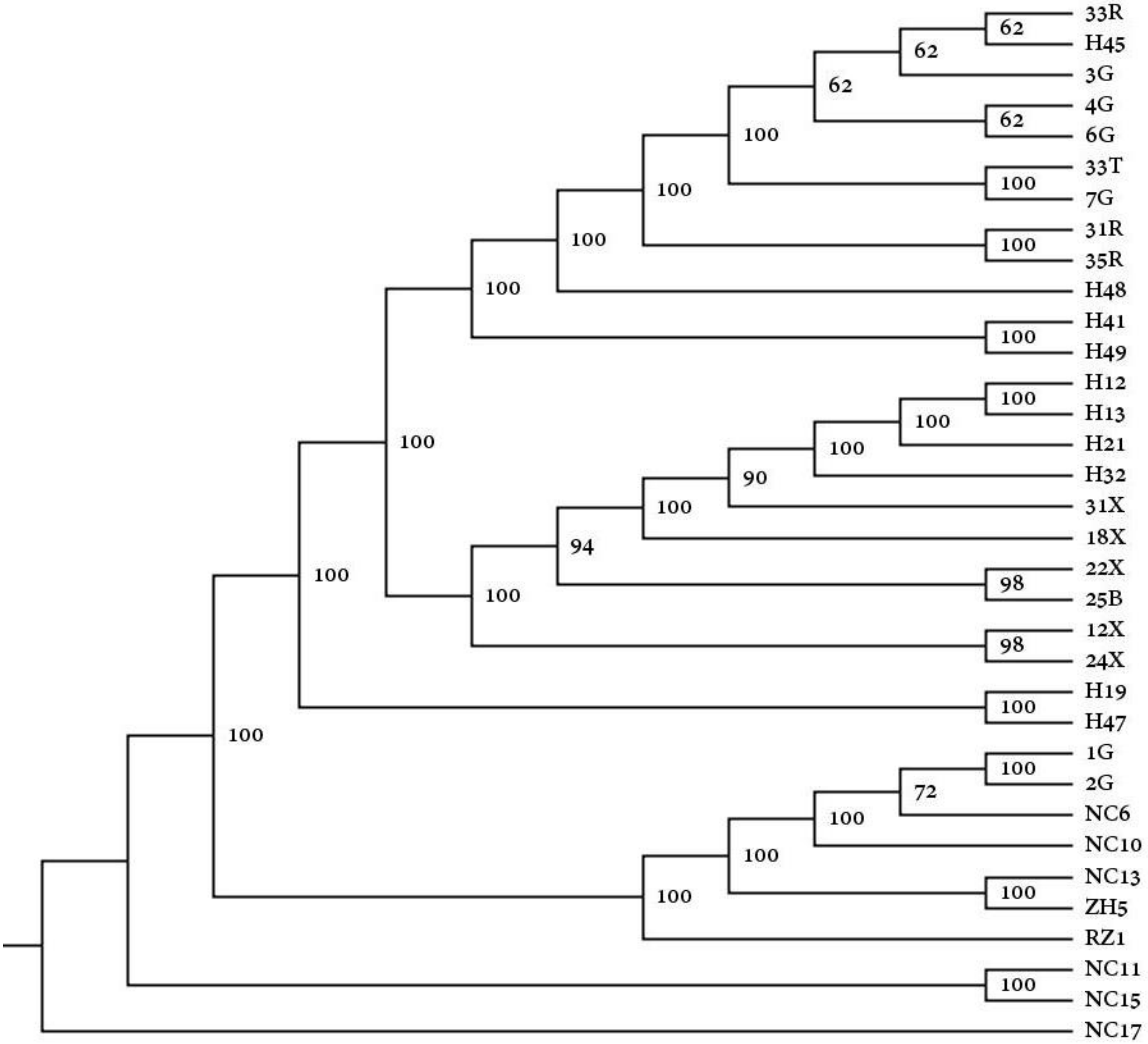

**Figure S4:** Maximum Likelihood cladogram based on 1540 nuclear genes corresponding with the N2 data set (Table 2), generated using IQtree. Bootstrap support values based on 1000 ultrafast bootstrap replicates are indicated at each node. Codes for taxon names are in Table 1.

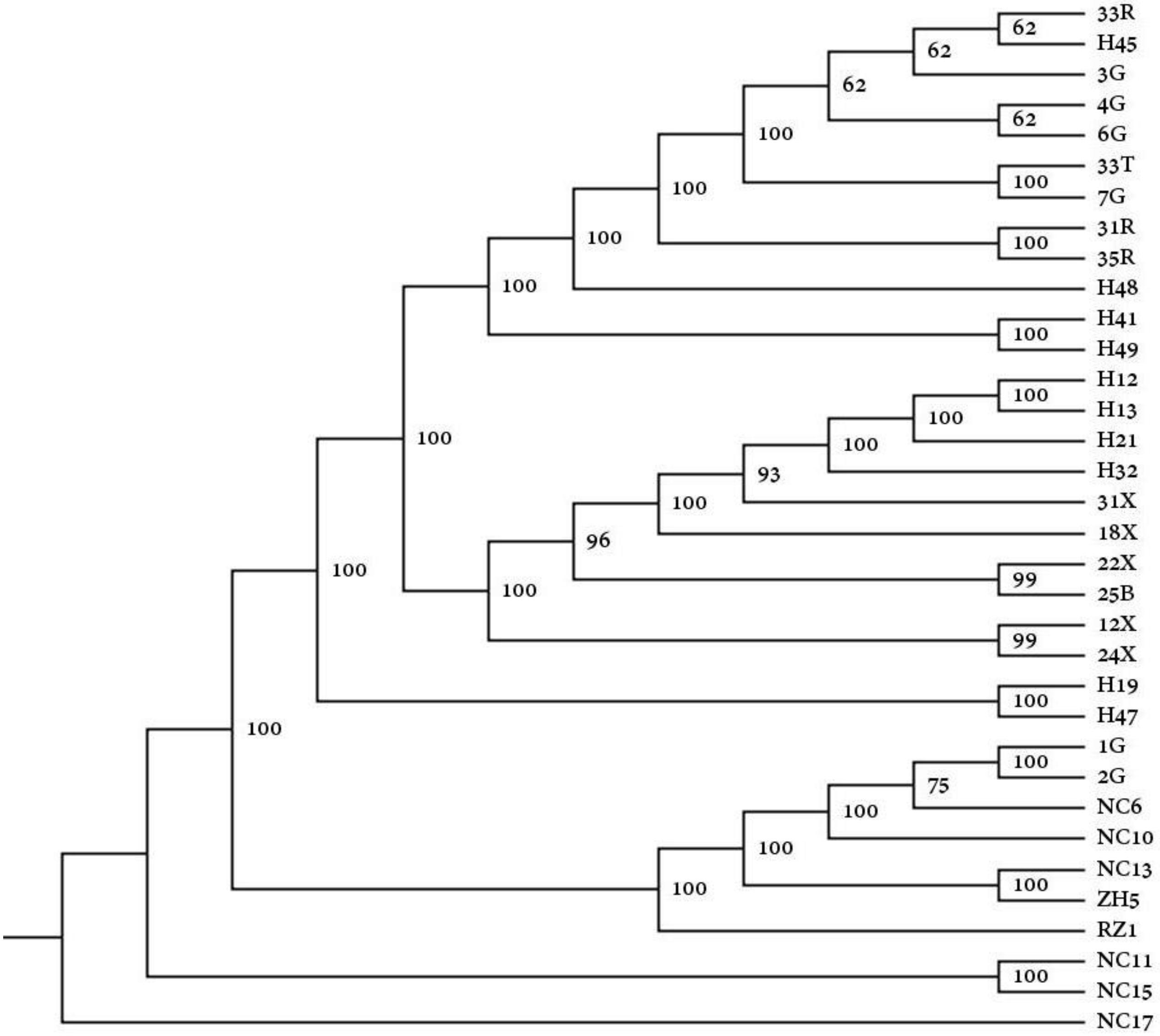

**Figure S5:** Maximum Likelihood cladogram based on 886 nuclear genes corresponding with the N4 data set (Table 2), generated using IQtree. Bootstrap support values based on 1000 ultrafast bootstrap replicates are indicated at each node. Codes for taxon names are in Table 1.

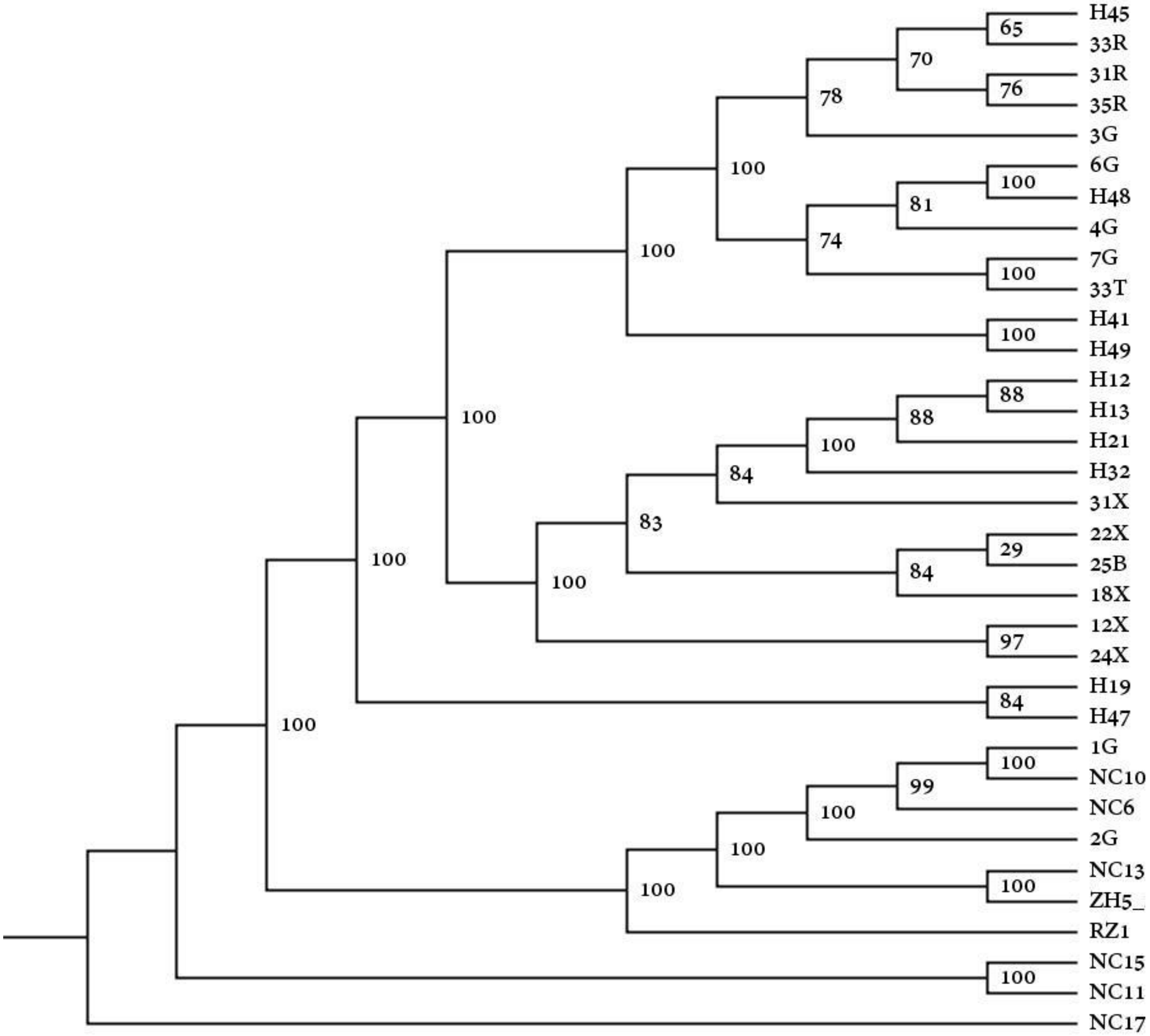

**Figure S6:** ASTRAL species tree based on 886 nuclear gene trees corresponding with the N5 data set (Table 2). Local posterior probability values at each branch. Codes for taxon names are in Table 1.

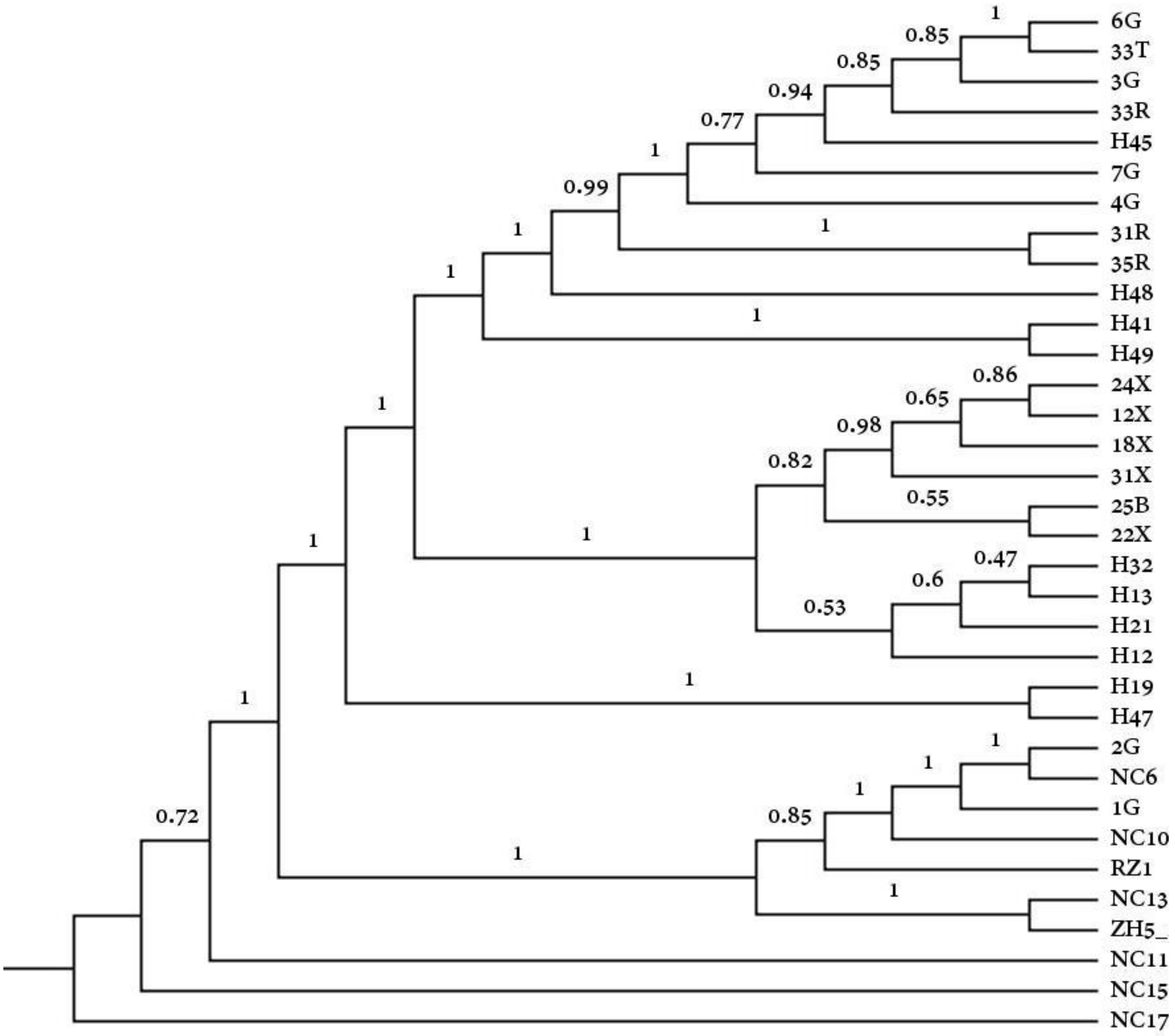

**Figure S7:** Maximum Likelihood cladogram based on complete plastomes corresponding with the CP1 data set (Table 2), generated using IQtree. Bootstrap support values based on 1000 ultrafast bootstrap replicates are indicated at each node. Codes for taxon names are in Table 1.

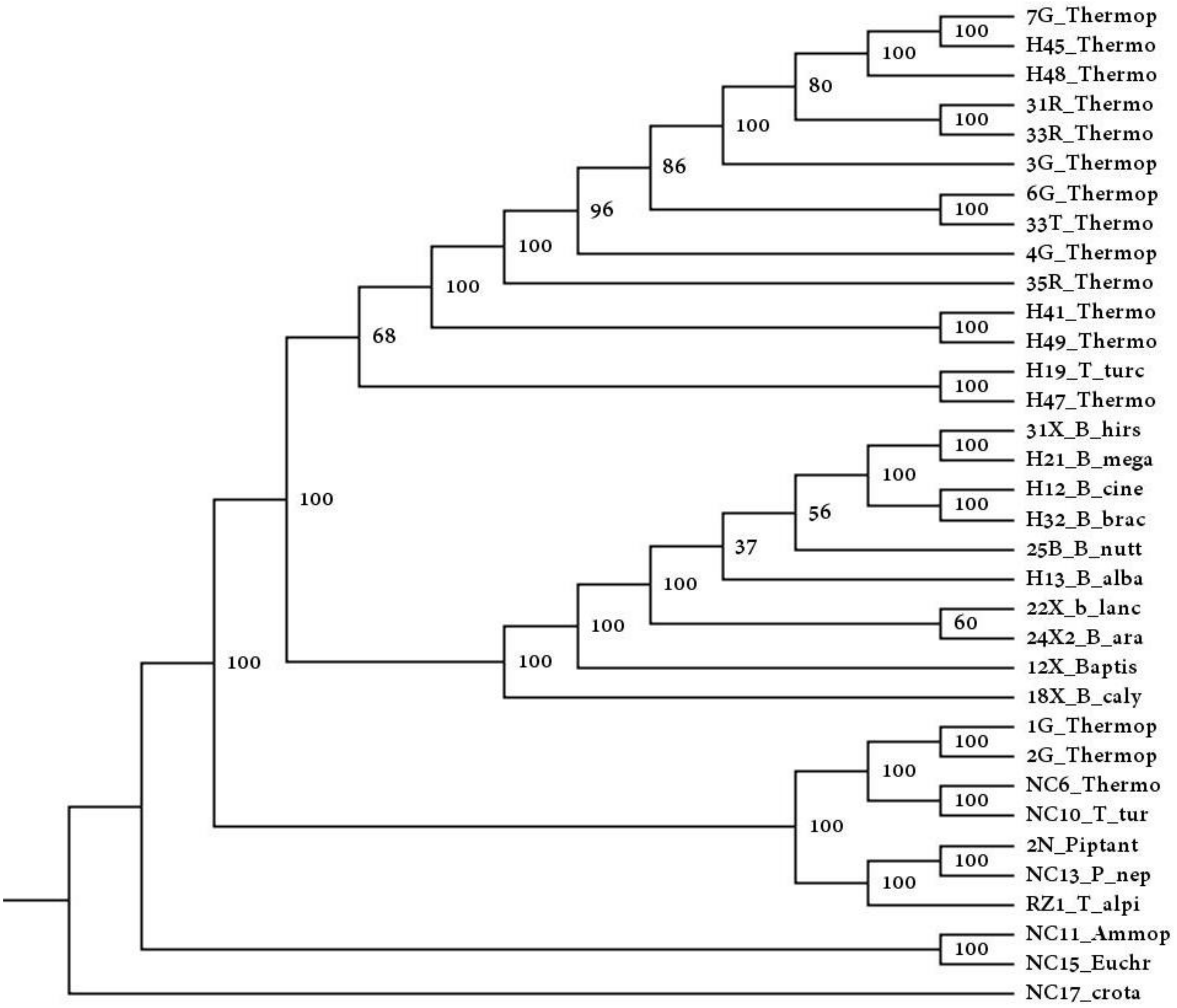

**Figure S8:** Maximum Likelihood cladogram based on complete plastomes corresponding with the CP2 data set (Table 2), generated using IQtree. Bootstrap support values based on 1000 ultrafast bootstrap replicates are indicated at each node. Codes for taxon names are in Table 1.

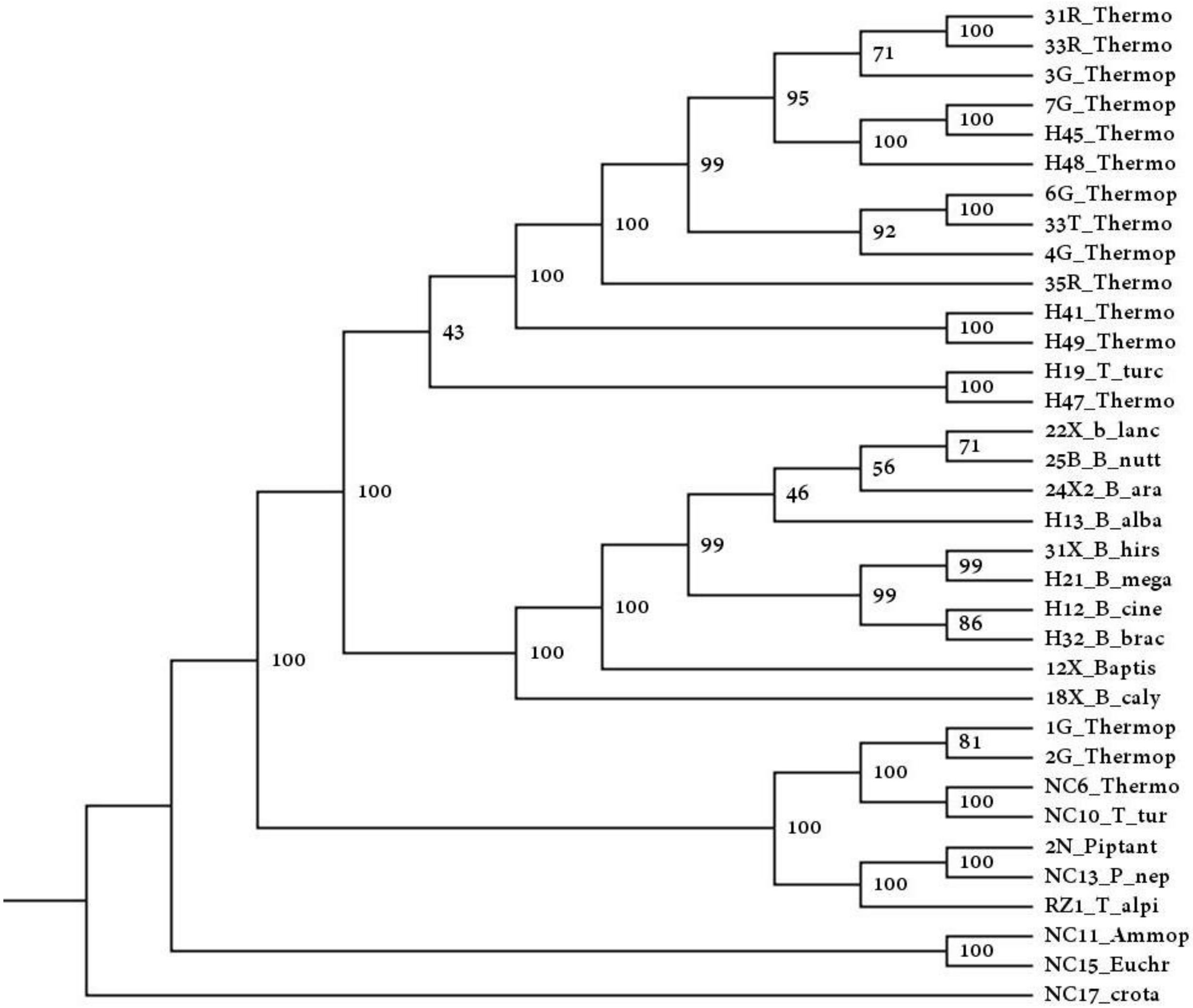

**Figure S9:** Maximum Likelihood cladogram based on complete plastomes corresponding with the CP3 data set (Table 2), generated using IQtree. Bootstrap support values based on 1000 ultrafast bootstrap replicates are indicated at each node. Codes for taxon names are in Table 1.

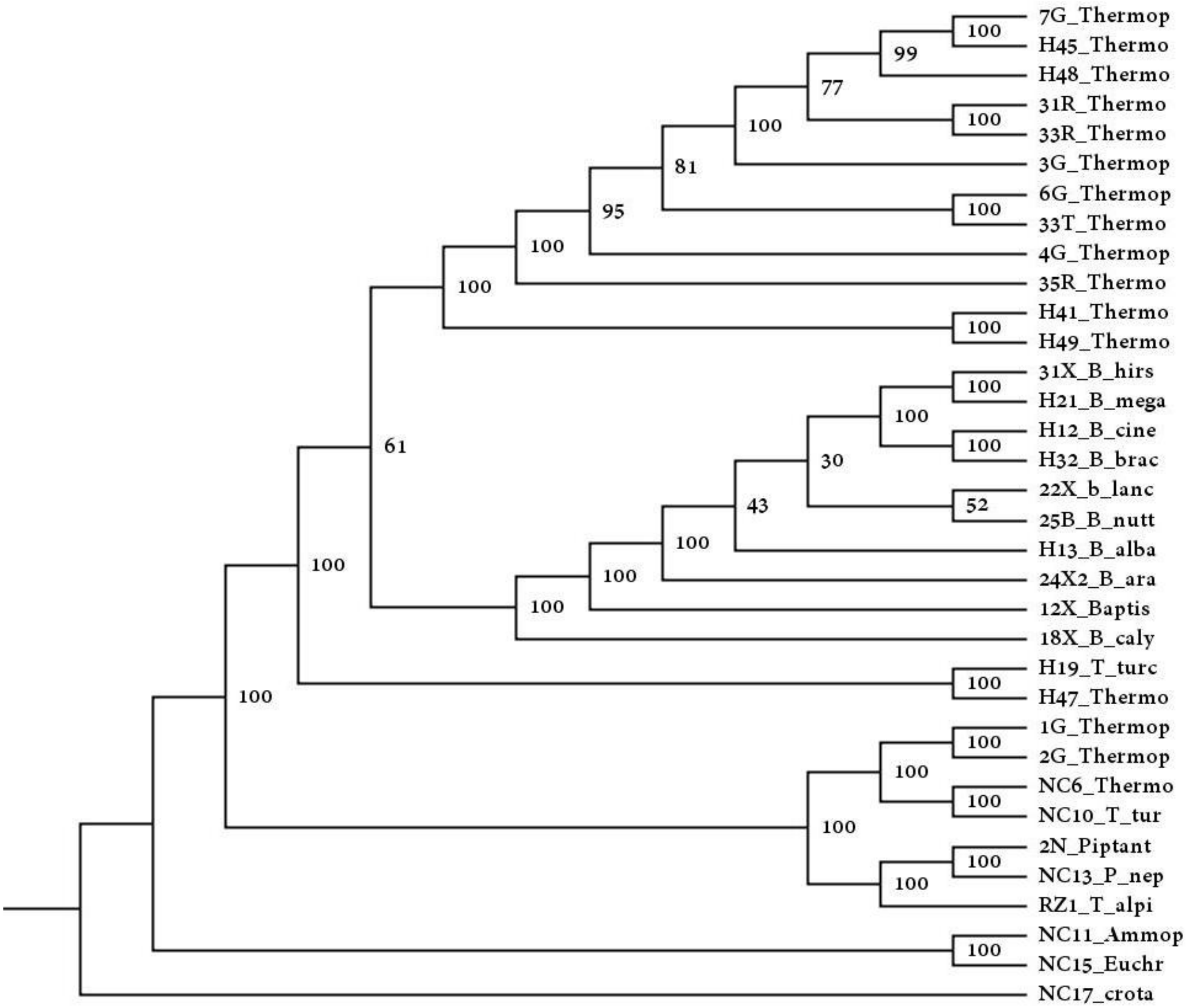

**Figure S10:** Maximum Likelihood cladogram based on complete plastomes corresponding with the CP4 data set (Table 2), generated using IQtree. Bootstrap support values based on 1000 ultrafast bootstrap replicates are indicated at each node. Codes for taxon names are in Table 1.

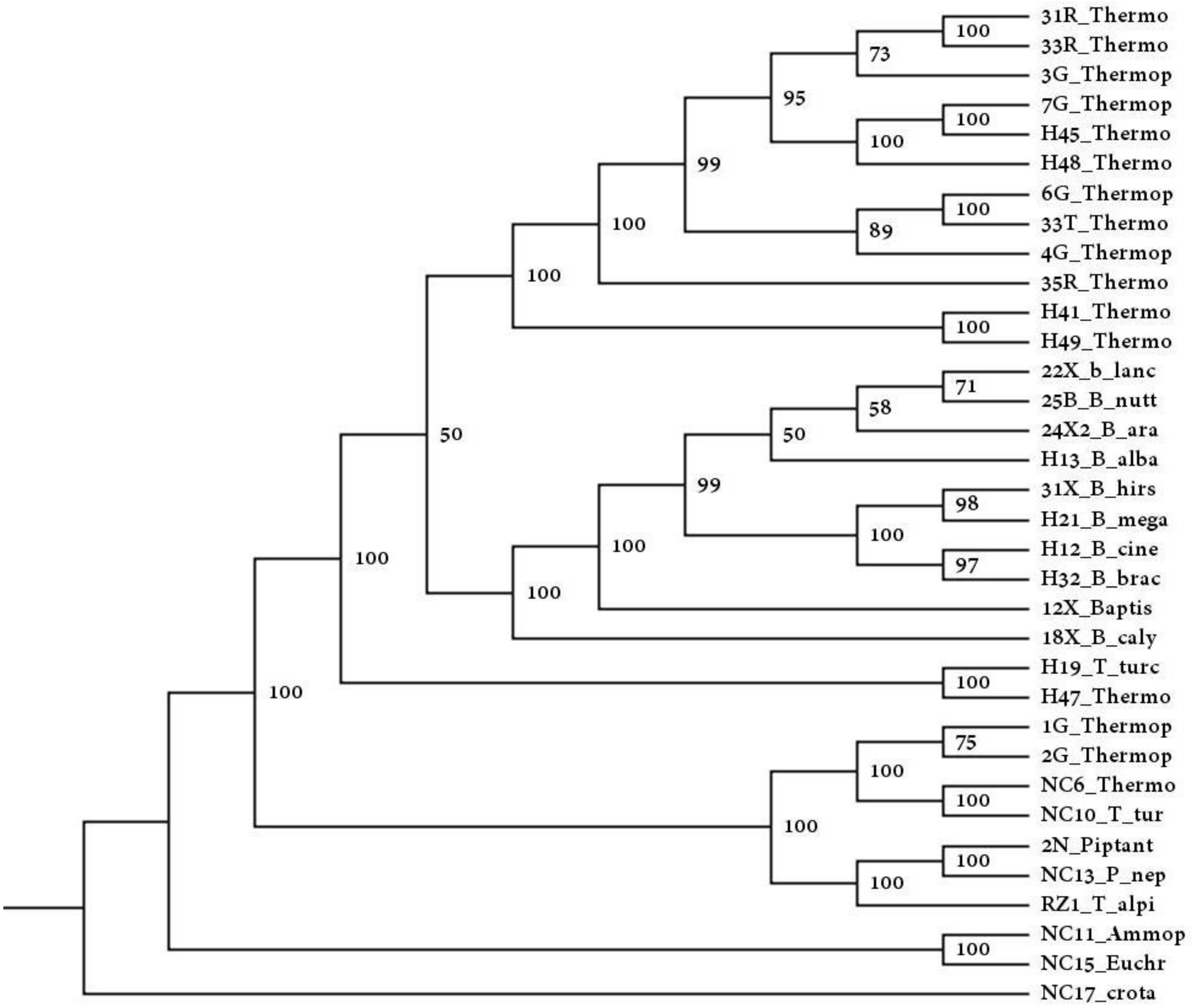

**Figure S11:** Maximum Likelihood cladogram based on 77 plastome CDS genes corresponding with the CP5 data set (Table 2), generated using IQtree. Bootstrap support values based on 1000 ultrafast bootstrap replicates are indicated at each node. Codes for taxon names are in Table 1.

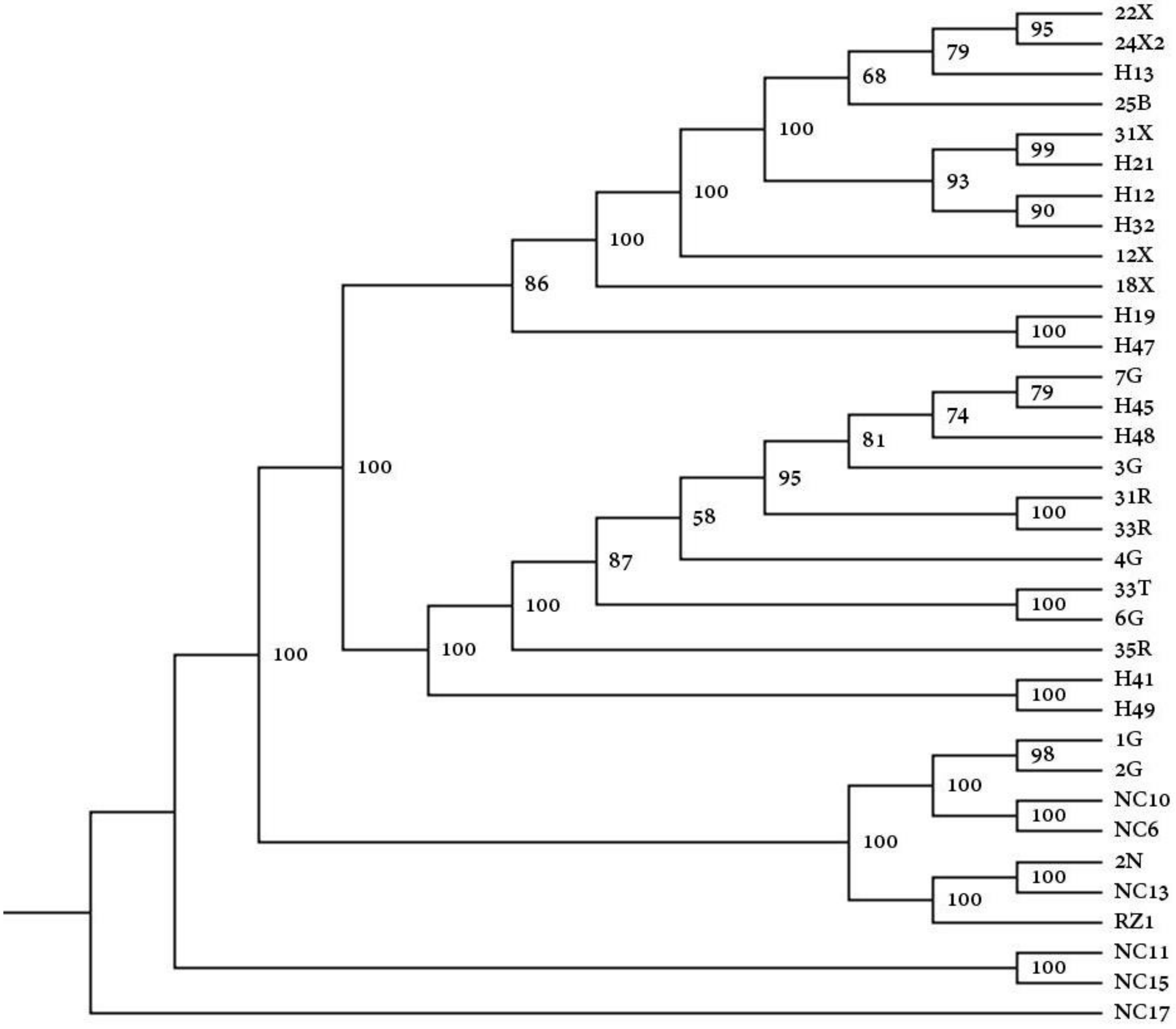

**Figure S12:** Maximum Likelihood cladogram based on 77 plastome CDS genes corresponding with the CP6 data set (Table 2), generated using IQtree. Bootstrap support values based on 1000 ultrafast bootstrap replicates are indicated at each node. Codes for taxon names are in Table 1.

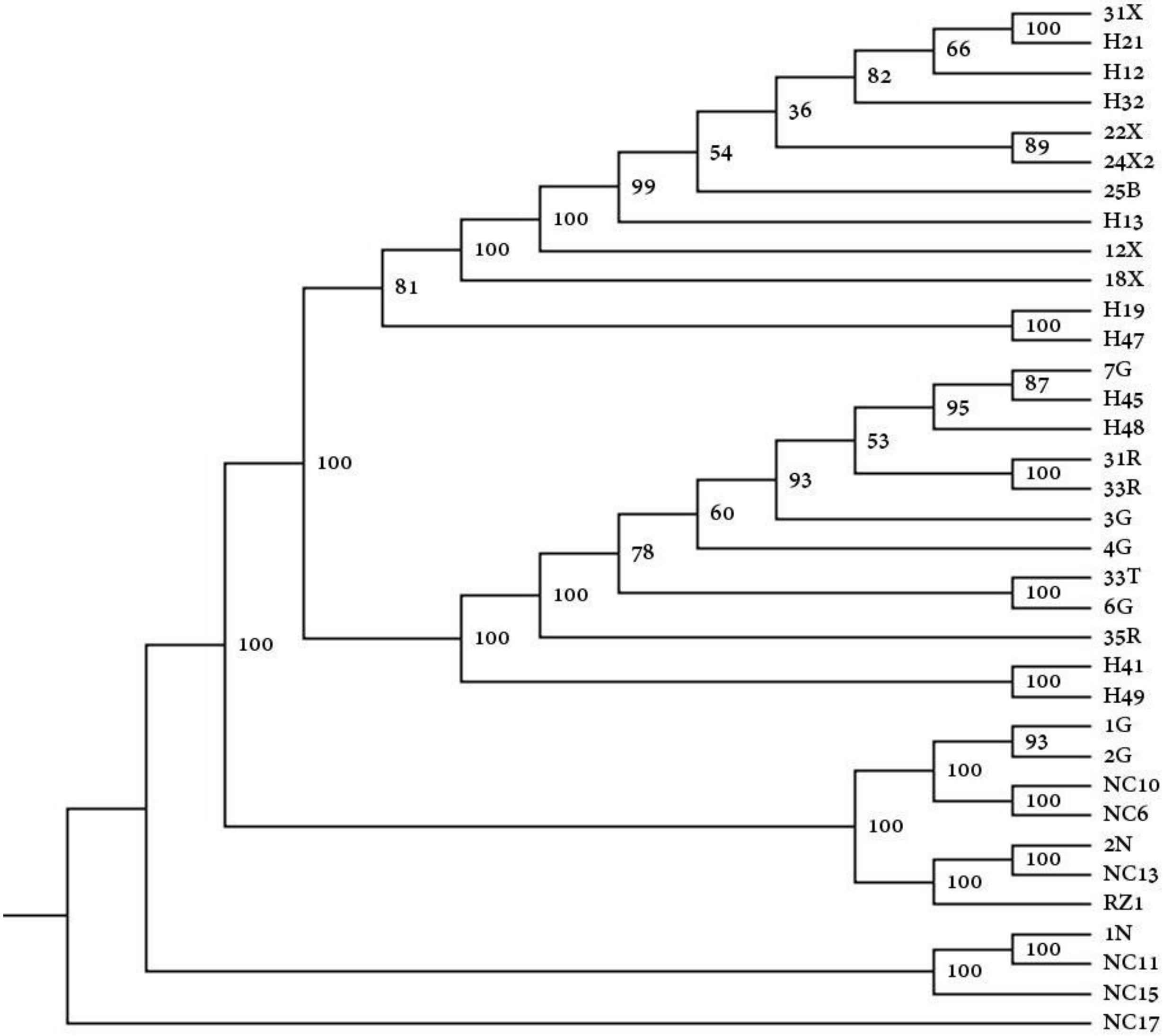

**Figure S13:** Maximum Likelihood cladogram based on 77 plastome CDS genes corresponding with the CP7 data set (Table 2), generated using IQtree. Bootstrap support values based on 1000 ultrafast bootstrap replicates are indicated at each node. Codes for taxon names are in Table 1.

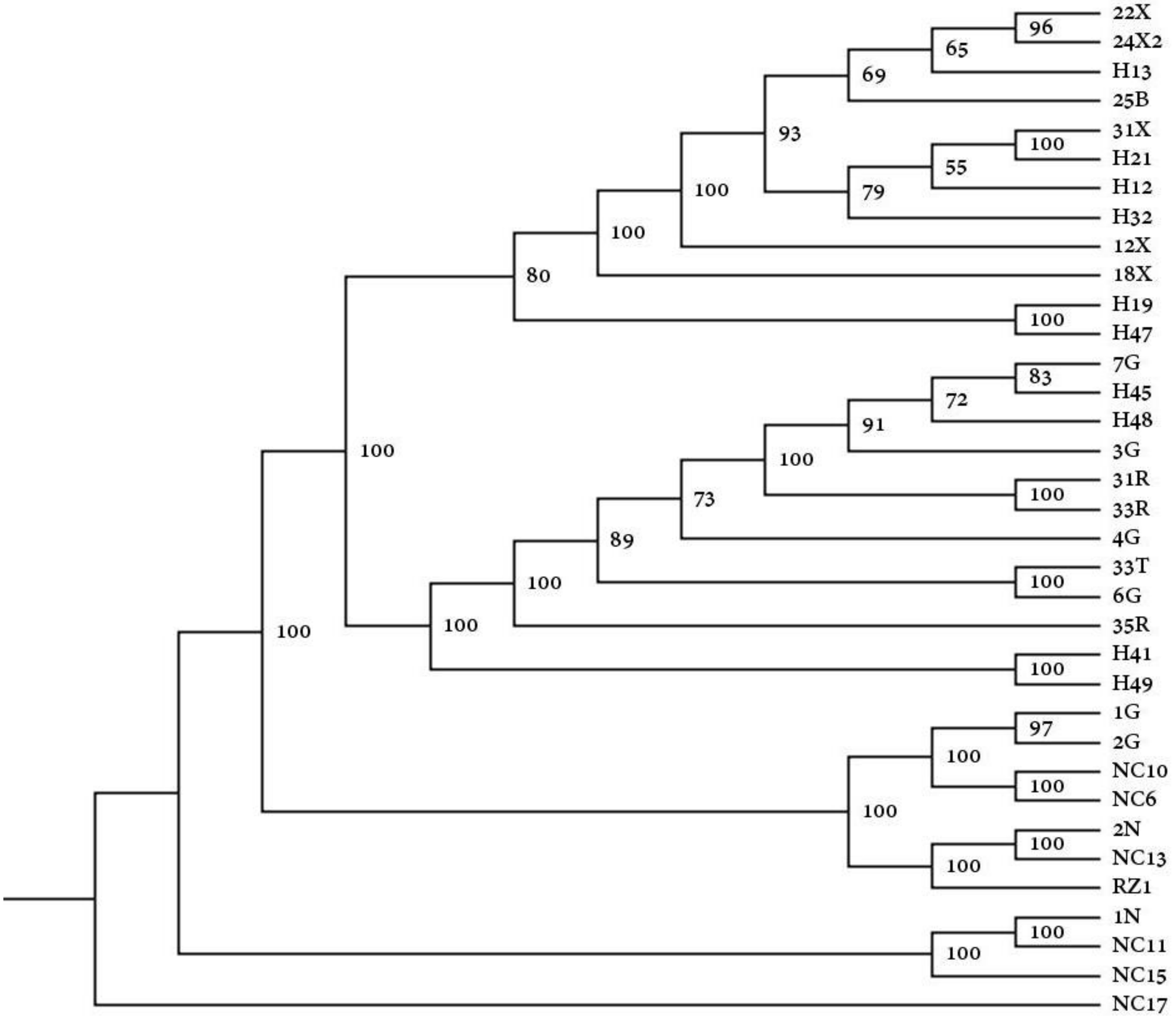

**Figure S14:** Maximum Likelihood cladogram based on 77 plastome CDS genes corresponding with the CP8 data set (Table 2), generated using IQtree. Bootstrap support values based on 1000 ultrafast bootstrap replicates are indicated at each node. Codes for taxon names are in Table 1.

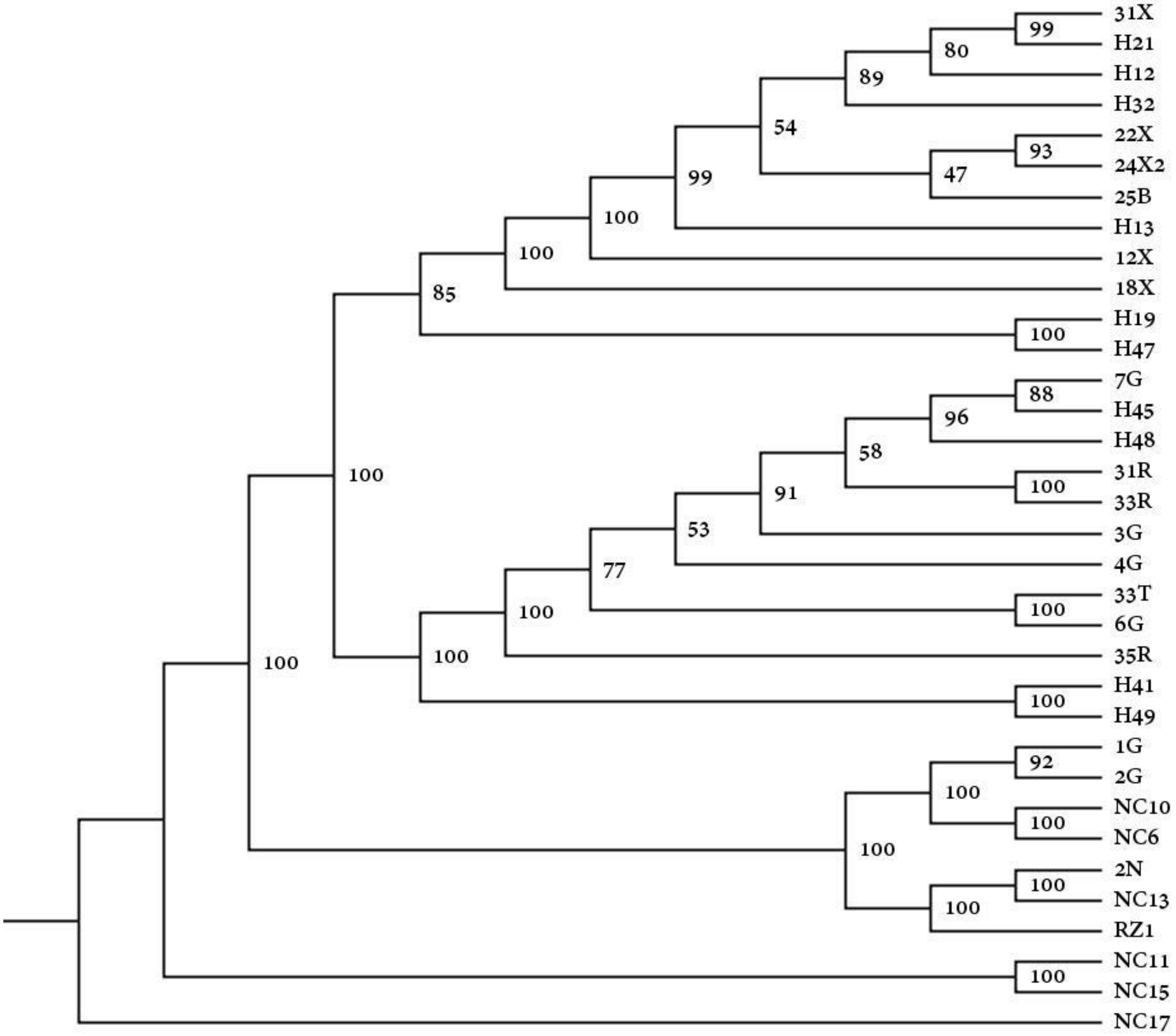

**Figure S15:** ASTRAL species tree based on 77 plastome CDS genes corresponding with the CP9 data set (Table 2). Local posterior probability values at each branch. Codes for taxon names are in Table 1.

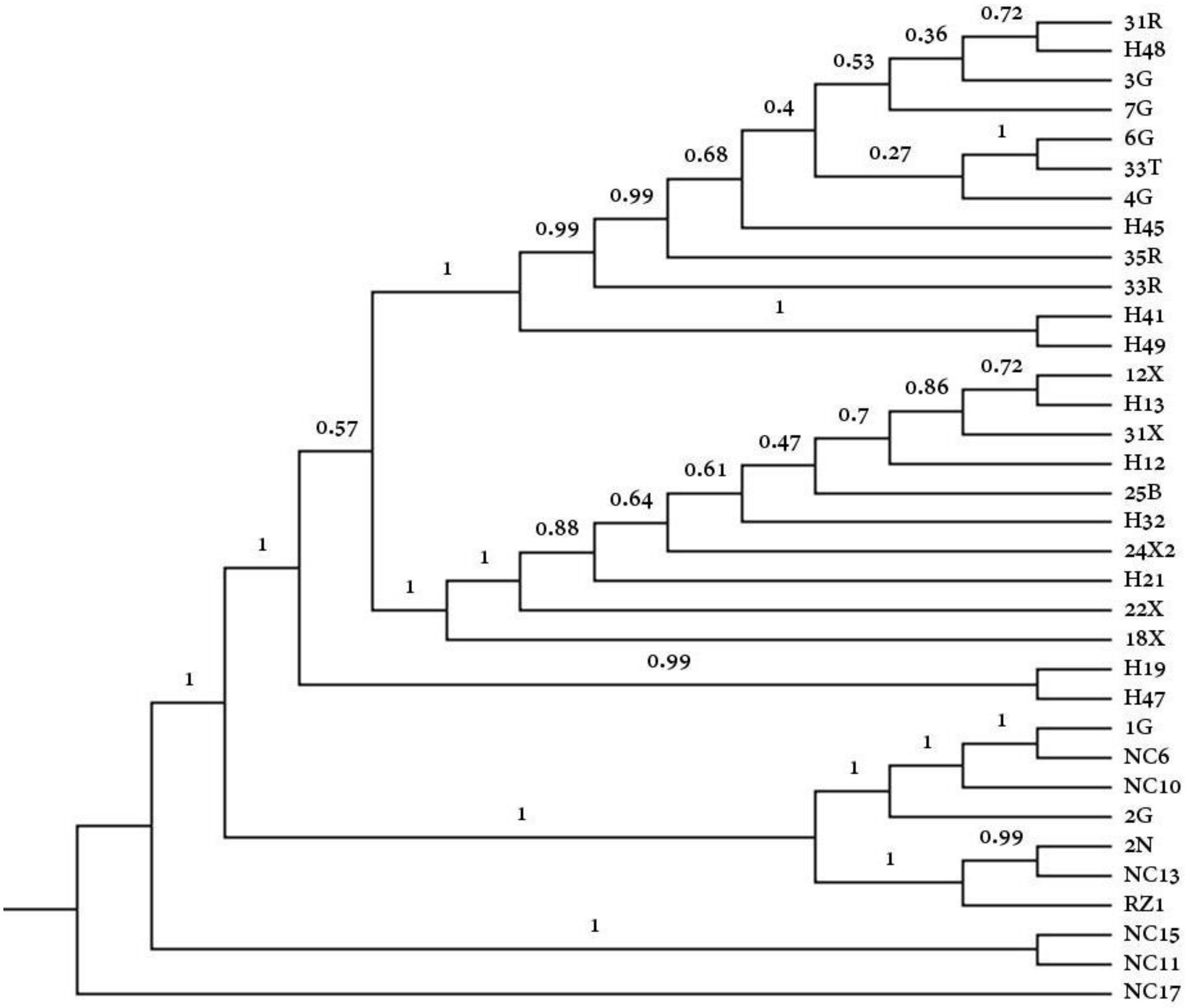
